# Altered GM1 catabolism affects NMDAR-mediated Ca^2+^ signaling at ER-PM junctions and increases synaptic spine formation

**DOI:** 10.1101/2023.07.10.548446

**Authors:** Jason A. Weesner, Ida Annunziata, Diantha van de Vlekkert, Camenzind G. Robinson, Yvan Campos, Ashutosh Mishra, Leigh E. Fremuth, Elida Gomero, Huimin Hu, Alessandra d’Azzo

## Abstract

Endoplasmic reticulum–plasma membrane (ER-PM) junctions mediate Ca^2+^ flux across neuronal membranes. The properties of these membrane contact sites are defined by their lipid content, but little attention has been given to glycosphingolipids (GSLs). Here, we show that GM1-ganglioside, an abundant GSL in neuronal membranes, is integral to ER-PM junctions; it interacts with synaptic proteins/receptors and regulates Ca^2+^ signaling. In a model of the neurodegenerative lysosomal storage disease, GM1-gangliosidosis, pathogenic accumulation of GM1 at ER-PM junctions due to β-galactosidase deficiency drastically alters neuronal Ca^2+^ homeostasis. Mechanistically, we show that GM1 interacts with the phosphorylated NMDAR Ca^2+^ channel, thereby increasing Ca^2+^ flux, activating ERK signaling, and increasing the number of synaptic spines without increasing synaptic connectivity. Thus, GM1 clustering at ER-PM junctions alters synaptic plasticity and exacerbates the generalized neuronal cell death characteristic of GM1-gangliosidosis.

## Introduction

In eukaryotes, communication among intracellular organelles or with the cell’s exterior is a prerequisite for cells to adapt their dynamic metabolic needs and respond to diverse physiologic or pathologic cues [1]. One universal mode of communication is through membrane contact sites (MCSs), which represent areas of juxtaposition between organellar membranes and/or the plasma membrane (PM), maintaining an average distance of ~10-35 nm [2, 3]. MCSs form transiently and reversibly by tethering membrane proteins or protein complexes and lipids, creating specialized “hit and run” signaling platforms that coordinate the synthesis/transport of lipids, the exchange of ions and metabolites (e.g., Ca^2+^), and other metabolic processes that do not require membrane fusion or vesicular trafficking [4–7]. The extensive membrane network of the endoplasmic reticulum (ER) makes contacts with virtually all other intracellular organelles and the PM [4, 8]. ER–mitochondria MCSs, named mitochondria associated–ER membranes (MAMs) were one of the first of these structures identified in electron micrographs of rat liver cells [9] and later studied at the biochemical level in relation to phospholipid biosynthesis and transfer, Ca^2+^ signaling, ER-mitochondria–mediated apoptosis, autophagy, mitochondrial dynamics, and energy metabolism [7, 10–14].

Many functions assigned to MAMs are shared by another type of contact site formed by apposition of the ER with the PM, referred to as ER-PM junctions [15]. These MCSs, which were first visualized on electron micrographs of muscle cells [16], frequently occur in neurons, where the highly polarized membranes may require ER-PM junctions to enable efficient synaptic communication [17–19]. Notably, a comprehensive analysis of the ER distribution in mouse brain tissue revealed that ER cisternae extend throughout the neuronal cell body, axons, and dendritic arbors, and they penetrate both the postsynaptic dendritic spines and the presynaptic axon terminals [18]. The quantity and type of synapses depend on the size, shape, and number of the dendritic spines, which are highly plastic and subjected to changes in response to neuronal activity [20, 21]. Two major classes of spines, large mushroom-shaped spines and thin-shaped spines, are controlled by the quantity and distribution of two glutamate receptors, N-methyl D-aspartate receptors (NMDARs) and α-amino-3-hydroxy-5-methyl-4-isoxazolepropionate receptors (AMPARs), that are integral to neuroplasticity at excitatory synapses [21–23]. Both receptors serve as ion channels, and their recruitment to or removal from the PM depends on their phosphorylation status and changes in Ca^2+^ flux at the postsynaptic terminal [24, 25]. The level, duration, and special distribution of Ca^2+^ influx through NMDARs can differentially activate the Ca^2+^-mediated kinase CaMKII or the Ca^2+^-mediated phosphatase calcineurin, which antagonistically control the shape and size of the spines [21, 22]. Although not yet demonstrated, the fact that the ER is differentially distributed in the spines based on their size/shape alludes to the formation of ER-PM junctions at synapses [18].

Like MAMs, ER-PM junctions are essential membrane hubs with two primary roles: they regulate Ca^2+^ flux, mostly in response to changes in ER Ca^2+^ levels, and they mediate membrane remodeling by transferring/shuttling lipids between the ER and the PM [26–28]. Canonical protein– protein interactions at ER-PM junctions are those between the ER-resident Ca^2+^ sensor, stromal-interaction molecule (STIM), and the PM-localized Ca^2+^ channel, ORAI, which promotes Ca^2+^ influx via the store-operated Ca^2+^-entry pathway, following Ca^2+^ release from the ER [29, 30]. Additional constituents of the ER-PM junctions, some of which are *bona fide* molecular tethers, include the vesicle-associated membrane protein–associated proteins (VAPs), phospholipid- and glycerolipid-binding/transport proteins, Ca^2+^ sensors, ion channels/receptors, and other membrane proteins whose structural and functional characteristics define the cellular processes that occur within these microdomains [15, 26, 27]. Equally relevant are the lipid components of ER-PM junctions, i.e., phospholipids and phosphoinositides, cholesterol, sphingomyelin, and glycosphingolipids (GSLs), because by influencing membrane fluidity, they facilitate the recruitment of specific proteins/protein complexes, thereby modulating their localized activities [15, 28, 31, 32]. In neurons, it is proposed that ER-resident phosphoinositides (PIs) are transported to the PM by transmembrane protein 24 (TMEM24), which re-localizes from the ER to the ER-PM junctions in response to Ca^2+^ signaling [33, 34]. PIs are converted into potent signaling molecules, such as phosphatidylinositol-4-phosphate (PIP) and phosphatidylinositol 4,5-bisphosphate (PIP2), at the PM via the activity of membrane-bound neuronal phosphatases and kinases [35, 36]. Altering the lipid profiles of MCSs contributes to neuropathogenesis in various neurodegenerative diseases [8, 37]. Until now, most studies have focused on the involvement of phospholipids and cholesterol, while less attention has been given to GSLs, particularly gangliosides [8].

Gangliosides are the most prominent sialoglycolipids in the mammalian brain [38]. The levels and expression pattern of gangliosides change in a regulated, region-specific manner during neuronal processes (e.g., neurogenesis, neuronal differentiation, axon and dendrite formation, synaptogenesis, and neurotransmission) [39]. Thus, maintaining their homeostatic concentration ensures the integrity and proper composition of neuronal membranes. Among the complex gangliosides, the monosialylated GM1 is the most abundant [40]. Because of its amphiphilic structure, GM1 can engage in hydrophobic and hydrophilic interactions with membrane receptors, ion (Ca^2+^) channels/transporters, or other extracellular ligands [40, 41]. In addition, GM1 can cluster transiently within the bilayer and help form and stabilize membrane subdomains, such as the lipid rafts or glycosphingolipid-enriched microdomains (GEMs), which are structural constituents of MCSs [41–44]. Within these membrane microdomains, GM1 has both signaling and regulatory functions that depend on its capacity to modulate intracellular Ca^2+^ homeostasis by directly binding Ca^2+^ ions or interacting with Ca^2+^ channels/pumps/transporters and/or Ca^2+^-binding proteins/receptors [40, 45, 46]. In neurons, changes in GM1 levels occur concomitantly to fluctuations in intracellular Ca^2+^, affecting neuritogenesis, spine formation, long term potentiation (LTP) and depression, and synaptic activity [40, 41]. For example, during neuritogenesis, GM1 stimulates neurite formation by interacting with the active phosphorylated forms of the neurotrophic receptors, tropomyosin receptor kinase (TrkA, B, and C). In addition, exogenously added GM1 increases the levels of brain-derived neurotrophic factor (BDNF), the cognate ligand of TrkB [47, 48]. GM1 can also alter synaptic architecture by sequestering AMPARs to the post-synaptic GEMs, preventing these receptors from binding to the AMPAR-trafficking complex that removes them from the PM [49].

Catabolism of GM1 occurs in lysosomes and is solely mediated by the lysosomal enzyme β-galactosidase (β-GAL) [50, 51]. Genetic mutations resulting in deficient β-GAL activity cause GM1-gangliosidosis, a severe neurodegenerative lysosomal storage disease affecting children, adolescents, and adults [50, 51]. The neuropathologic course of GM1-gangliosidosis has been largely attributed to deregulation of cellular pathways triggered by the accumulation of GM1 in neurons [50, 51]. These features are fully replicated in *β-Gal^−/−^* mice that develop severe neurologic abnormalities early in life (i.e., tremor, ataxia, and paralysis of the hind limbs) that are accompanied by accumulation of GM1 in neurons of the brain and spinal cord, leading to massive cell death and neuroinflammation [52, 53]. At the molecular level, we have shown that GM1 accumulates in the GEM fraction of MAMs, where it binds to the phosphorylated/active form of the ER-resident inositol trisphosphate receptor (IP3R-1) Ca^2+^ channel, driving Ca^2+^ efflux from the ER into the mitochondria through a tethering Ca^2+^ megapore [42]. This elicits the simultaneous activation of the unfolded protein response and mitochondria-mediated apoptosis [42, 54]. However, the pathogenic consequences of altered GM1 concentration at the PM, where the ganglioside normally resides, have not been explored.

Here we show that GM1 not only is an integral component of ER-PM junctions but also favors the tethering of the apposed membranes. Increased levels of GM1 in *β-Gal^−/−^* neurons is accompanied by increased formation of ER-PM junctions, which affect overall synaptic architecture. We propose a mechanism by which active phosphorylated NMDARs (pNMDARs) are retained at *β-Gal^−/−^* ER-PM junctions via interactions with GM1. This in turn activates Ca^2+^-mediated extracellular signal–regulated kinase (ERK) signaling, thereby increasing dendritic spine formation without affecting synaptic density.

## Results

### *β-Gal^−/−^* Purkinje cells have increased formation of ER-PM junctions

To investigate the effects of GM1 accumulation on the formation, number, and structural features of ER-PM junctions, we first visualized these MCSs at the ultrastructural level by using transmission electron microscopy (TEM). We analyzed Purkinje cells because of their large cell bodies and extensive pathology in *β-Gal^−/−^* mice, as previously reported [52, 55]. Morphologic comparison of wild-type (WT) and *β-Gal^−/−^* Purkinje cells revealed that a high proportion of the ER membranes was juxtaposed to the PM in the *β-Gal^−/−^* cells, forming a nearly continuous network of ER-PM junctions (**Figure 1A, yellow lines**). This distribution differed from that in WT cells, where the MCSs appeared dispersed, fragmented, and less numerous (**Figure 1A, yellow lines**). The extensive formation of ER-PM junctions in *β-Gal^−/−^* Purkinje cells, compared to WT cells, was also confirmed quantitatively by calculating the percentage of tethering events between the ER and the PM in WT mice and *β-Gal^−/−^* mice at 1-, 3-, and 6-months of age (**Figure 1B)**.

**Figure 1:**
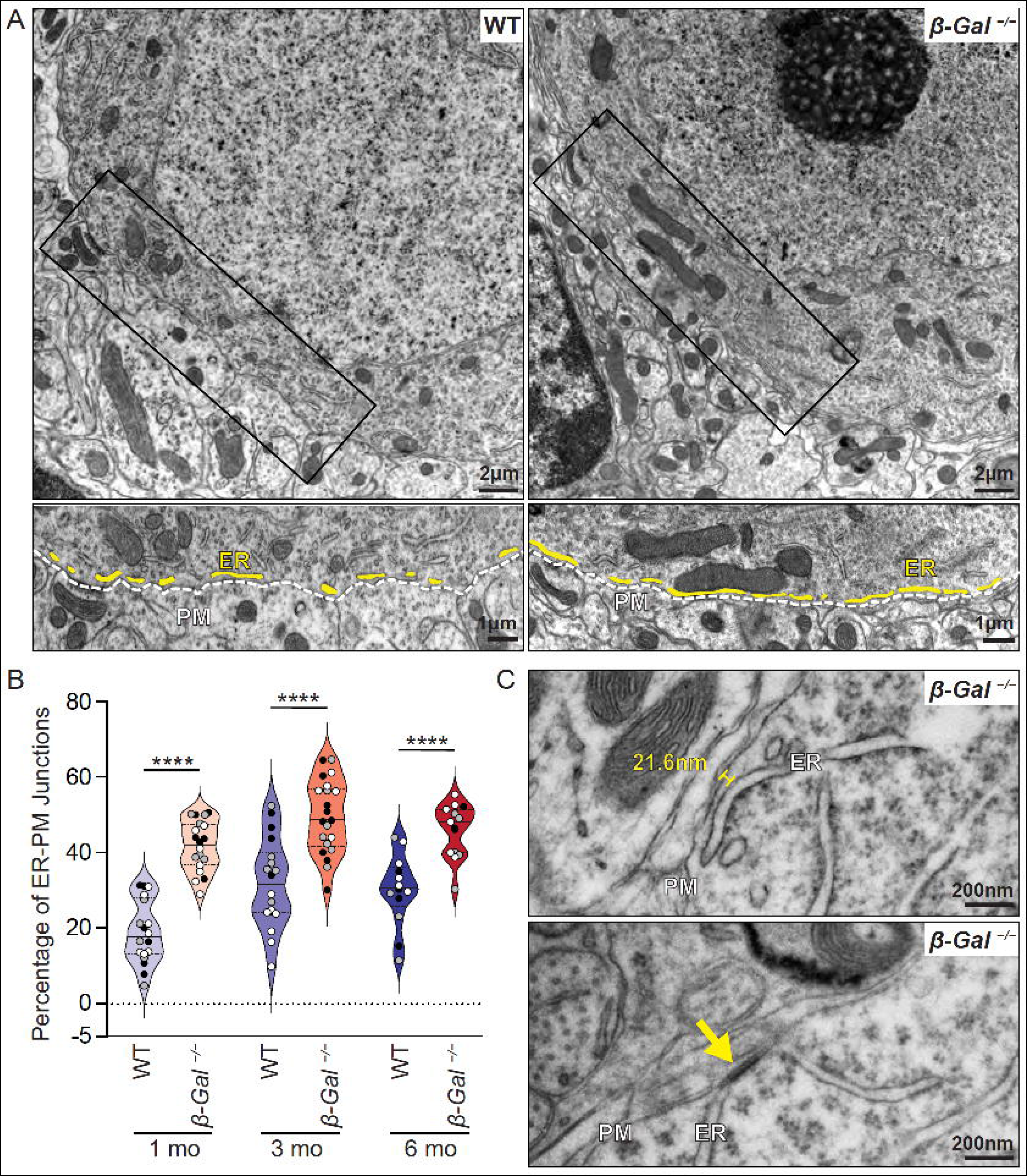
Increased formation of ER-PM junctions in *β-Gal^−/−^* Purkinje cells. (A) TEM images of Purkinje cells in 6-month-old WT and *β-Gal^−/−^* mice. Scale bar: 2 µm. Bottom images show 1.4× zoom of black boxes with highlighted PM (white line) and ER fragments (yellow lines). Scale bar: 1 µm. (B) Quantification of the average area of ER tethered to the PM (percentage) in 1-, 3-, and 6-month-old WT and *β-Gal^−/−^* mice. *n* = 3 mice; 5-9 Purkinje cells per mouse, each mouse is represented by a different colored filled datapoint (black, grey, white). Values are expressed as mean ± SD. Statistical analysis was performed using the One-way ANOVA; \*\*\*\**p* < 0.0001. (C) High-magnification TEM images of Purkinje cells from a 6-month-old *β-Gal^−/−^* mouse shows ER-PM junctions that are defined by the distance between the membranes: the top image shows separation by a distance ≤35 nm, and the bottom image shows close apposition resulting in a density (yellow arrow). Scale bar: 200 nm. ER: endoplasmic reticulum; PM: plasma membrane.

A significant increase in the percentage of ER-PM junctions was observed in the *β-Gal^−/−^* Purkinje cells compared to that in the WT cells, suggesting that the increased concentration of GM1 at both membranes favors the formation of these MCSs (**Figure 1B**). In both WT and *β-Gal^−/−^* Purkinje cells, the average distance between the apposing membranes was <35 nm (**Figure 1C, top image and S1A, left image**), as previously reported in other cell types [2, 3]. Occasionally, portions of the ER cisternae involved in junction formation narrowed into high-density regions, usually predictive of high protein content, where the ER lumen collapsed and the space between the ER membranes and the juxtaposed PM was no longer visible (**Figure 1C, bottom image and S1A, left image; yellow arrows**). These electron-dense regions were also present in unusual triple-contact sites formed by tethering mitochondria to ER-PM junctions (**Figure S1B**). These triple MCSs were rarely seen in WT samples but were numerous and readily detected in the *β-Gal^−/−^* cells.

Another feature of contacts that the ER forms with the PM or other organellar membranes was the repositioning of ribosomes at the contact sites. Ribosomes appeared to be excluded from the face of the cisternae closest to the PM, a configuration that facilitates the tight association of ER membranes at MCSs (**Figure 1C, top image**). These different structural configurations of ER-PM junctions, indicative of their functional status, were maintained in both WT and *β-Gal^−/−^* neurons; however, the increase in ER-PM junction formation in *β-Gal^−/−^* Purkinje cells suggests changes in the cellular pathways that control these MCSs.

### Exogenous supplementation of GM1 promotes the formation ER-PM junctions in WT cells

We next determined whether the increased number of ER-PM junctions was directly linked to GM1 accumulation at the respective membranes. We used HeLa cells exogenously loaded with GM1 and transduced with the GFP-expressing construct MAPPER [56] that selectively detects the formation of ER-PM junctions in real-time, along the PM surface, in response to Ca^2+^ depletion from the ER.

At baseline, GFP^+^ puncta had already formed along the cell surface of GM1-loaded cells but were hardly detectable in untreated cells (**Figure 2A**). Live-cell imaging of untreated and GM1-loaded cells for up to 10 min revealed a significant increase in GFP^+^ puncta in response to thapsigargin-induced Ca^2+^ depletion from the ER (**Figure 2A**). GFP^+^ puncta begun to build up along the PM at 1.5 min after thapsigargin administration; after 3 min, the fluorescence intensity was significantly higher at the cell surface of GM1-loaded cells (**Figure 2A, B**). The GM1-loaded cells also had GFP^+^ puncta along the entire PM, but control cells showed a limited number of contact sites (**Figure 2A**), demonstrating a direct correlation between the level of GM1 and ER-PM junction formation, which was potentiated by the release of Ca^2+^ from the ER. After 6 min, GFP fluorescence in untreated HeLa cells returned to baseline, while GM1-loaded cells expressed GFP throughout the experiment. No response was measured after dimethylsulfoxide (DMSO) treatment (**Figure 2B**).

**Figure 2:**
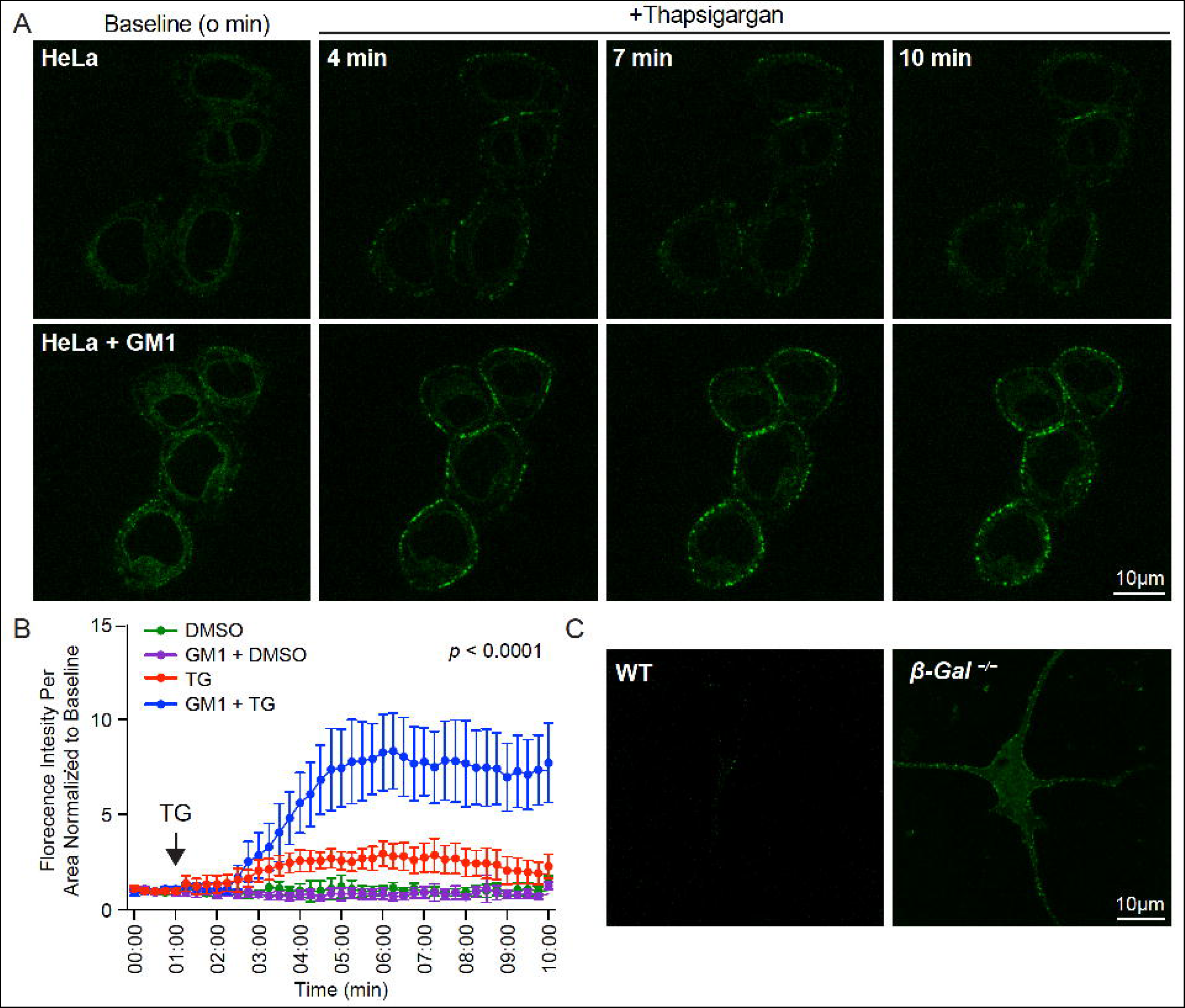
ER-PM junctions form in GM1-loaded HeLa cells and primary neurons. (A) Time-course confocal immunofluorescence of MAPPER-transduced HeLa cells (green). GM1 was exogenously added to HeLa cells. Images were captured at baseline (0 min) and at 4-, 7-, and 10-min post treatment with thapsigargin (TG). (B) Quantification of fluorescence intensity per area over time (0–10 min). Experiments were performed on HeLa cells in triplicate, and 6-7 fields of view were analyzed; DMSO: *n* = 19, GM1 + DMSO: *n* = 20, TG: *n* = 19, and GM1 + TG: *n* = 17. Values are expressed as mean ± SD. Statistical analysis was performed using the Two-Way ANOVA tests; *p* < 0.0001. (C) Confocal images of MAPPER-transduced primary neurons isolated from P5 WT and *β-Gal^−/−^* pups show soma and neurite projections. Green puncta along the cell surface indicate ER-PM junctions. Scale bar: 10 µm.

A similar experiment conducted in *β-Gal^−/−^* primary neurons transduced with MAPPER showed GFP^+^ puncta along the entire PM of the soma and neuronal projections at baseline, but little-to-no puncta were visible in WT neurons (**Figure 2C**). These results unequivocally linked high levels of GM1 in neuronal membranes with increased formation of ER-PM junctions.

### GM1 accumulates in the ER-PM junctions isolated from *β-Gal^−/−^* brains

To identify all proteins and lipids present at the ER-PM junctions isolated from WT and *β-Gal^−/−^* brains, we optimized a purification method that enabled the enrichment of fractions containing ER, PM, and ER-PM junctions (**Figure S2A,B**). These fractions were tested for purity on immunoblots probed with the appropriate organelle markers. Proteins that populate ER-PM junctions, such as ORAI1, STIM1, STIM2, and VAP A and B isoforms (VAPA/B), were enriched in both WT and *β-Gal^−/−^* fractions, which also contained reduced levels of the ER and PM markers calnexin and N-cadherin, thereby confirming the suitability of this protocol for isolating these MCSs (**Figure 3A**). Moreover, anti-GM1 antibodies detected a massive accumulation of the ganglioside in all three *β-Gal^−/−^* fractions but especially in the ER and ER-PM junctions (**Figure 3A**). These results were further corroborated by high-performance thin-layer chromatography (HPTLC) assays (**Figure 3B**). The levels of GM1 were so high in the *β-Gal^−/−^* fractions that we needed to increase the concentration of the WT fractions by 3-fold to detect a signal. Band quantifications after normalization to loading from 7 independent experiments revealed that the highest GM1 levels were in the ER-PM junction fractions from WT and *β-Gal^−/−^* brains, indicating that GM1 is a normal constituent of these MCSs (**Figure 3C**). However, the amount of GM1 in the *β-Gal^−/−^* ER-PM junctions was almost 10 times greater than that in WT controls (**Figure 3C**).

**Figure 3:**
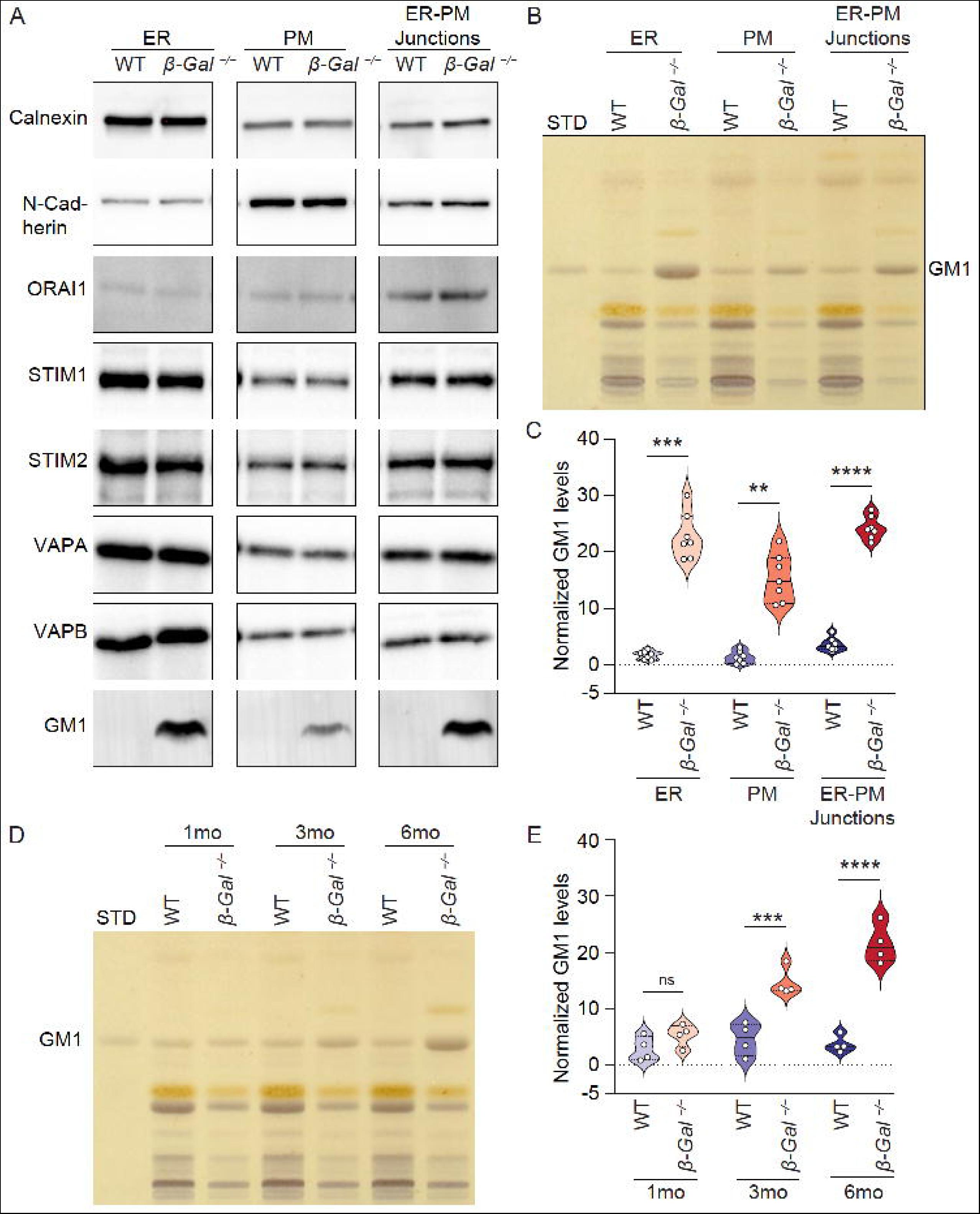
GM1 accumulates in *β-Gal^−/−^* ER-PM junctions. (A) Immunoblot analysis of cell fractions from 6-month-old WT and *β-Gal^−/−^* mice. Markers for the ER (Calnexin), PM (N-cadherin), and ER-PM junctions (ORAI1, STIM1, and STIM2, VAPA, and VAPB) were enriched in their respective fractions. Immunoblots using anti-GM1 antibody show high levels in *β-Gal^−/−^* fractions. (B) Representative HPTLC plate showing GM1 levels in the ER, PM, and ER-PM junctions isolated from 6-month-old WT and *β-Gal^−/−^* mice. STD: standard. Note: To detect GM1 in WT samples, the sample volume loaded was 3× that of the *β-Gal^−/−^* samples. (C) Quantification of GM1 levels from HPTLC plates performed in B. *n* = 7. Values are expressed as mean ± SD. Statistical analysis was performed using the Student’s *t*-test; \*\**p* < 0.01, \*\*\**p* < 0.001, \*\*\*\**p* < 0.0001. (D) Representative HPTLC plate showing GM1 levels in ER-PM junctions isolated from 1-, 3-, and 6-month-old WT and *β-Gal^−/−^* mice. To detect GM1 in WT samples, the sample volume loaded was 3× that of the *β-Gal^−/−^* samples. (E) Quantification of GM1 levels from HPTLC plates performed in D. *n* = 4. Values are expressed as mean ± SD. Statistical analysis was performed using the Student’s *t*-test with Welch’s correction; ns: not significant, \*\*\**p* < 0.001, \*\*\*\**p* < 0.0001.

Although the weights of the WT and *β-Gal^−/−^* brains were comparable at the start of the purification procedure, the fraction comprising the ER-PM junctions from the *β-Gal^−/−^* brains consistently contained a higher protein and lipid content than did the corresponding WT fraction (**Figure S2C**), further supporting the notion that *β-Gal^−/−^* neurons have more of these MCSs then do WT neurons. Using HPTLC, we monitored how GM1 levels changed in ER-PM junctions isolated from 1-, 3-, and 6-month-old WT and *β-Gal^−/−^* mouse brains (**Figure 3D**). GM1 bands were quantified in 4 independent assays (**Figure 3E**). The *β-Gal^−/−^* ER-PM junctions showed an accumulation of GM1 as the animals aged, with significantly higher levels of the ganglioside, compared to that in the WT fractions, starting at 3 months (**Figure 3E**). The GM1 levels in the WT ER-PM junctions remained similar at all ages, suggesting that GM1 gradually clusters at these MCSs only during disease progression.

### Comparative proteomic analysis of *β-Gal^−/−^* ER-PM junctions predicts upregulation of the synaptogenesis signaling pathway

We next employed a label-free, spectral-count comparative proteomic analysis to examine which proteins were differentially expressed in the ER-PM junctions isolated from WT and *β-Gal^−/−^* brains. The variance between the samples was low, and a total of 2212 proteins were identified (ProteomeChange:<u>PXD042994</u>), most of which resided in the ER/cytoplasm (1288), and the PM (513) (**Figure S3A**). The majority of the identified proteins belonged to major families of kinases, proteases, enzymes, transporters, ion channels, and G-protein–coupled receptors (ProteomeChange:<u>PXD042994</u>). Notably, among them were *bona fide* components of ER-PM junctions, including ORAI2 and TMEM24, both of which were clearly upregulated in the *β-Gal^−/−^* samples (**Figure S3B**).

The ER Ca^2+^-release channels IP3R-3 and ryanodine receptor types 2 and 3 (RYR2 and RYR3), were also increased (**Figure S3B**), which support the deregulated activity of the phosphorylated form of IP3R-1 observed in the *β-Gal^−/−^* MAMs [42]. Other proteins of interest that were differentially expressed in *β-Gal^−/−^* ER-PM junctions were the glutamate ionotropic receptor NMDAR-type 1 subunit, which increased 10-fold and TrkB, which increased 1.6-fold (**Figure S3B**). Both receptors are associated with LTP and promote postsynaptic spine formation by activating mitogen–activated protein kinase (MAPK)/ERK signaling pathway [47, 57]. We found that both ERK1 (a.k.a. MAPK3; 1.5-fold increase) and ERK2 (a.k.a. MAPK1; 1.3-fold increase) were increased in the *β-Gal^−/−^* samples (**Figure S3B**). In addition, the membrane-bound kinases, phosphatidylinositol 4-kinase type 2A (PI4K2A), phosphatidylinositol-4-phosphate 5-kinase type 1C (PIP5K1C), which phosphorylate PIs, showed some of the highest predicted fold increases (**Figure S3B**).

We further validated some of the proteins upregulated in the *β-Gal^−/−^* ER-PM junctions on immunoblots and quantified them by comparing WT and *β-Gal^−/−^* fractions (**Figure 4A-F and S3C-F**). The levels of NMDAR-type 1 subunit, TMEM24, PI4K, PIP5K1, and TRKB were significantly higher in the *β-Gal^−/−^* ER-PM junctions (**Figure 4A-F**). Given that the activated form of NMDAR-type 1 subunit, which is phosphorylated at Ser896 (pNMDAR), traffics the receptor to the PM and accumulates at the synapses [58], we also tested its levels and found it to be significantly increased in *β-Gal^−/−^* ER-PM junctions (**Figure 4A, S3C**). Although ERK1/2 levels did not differ between WT and *β-Gal^−/−^* fractions, the active phosphorylated forms of ERK1/2 (pERK1/2) were significantly increased in the *β-Gal^−/−^* ER-PM junctions (**Figure 4B**). Moreover, the increased ratio of pERK/ERK inferred enhanced ERK signaling at these MCSs (**Figure S3D**). We found that BDNF, the cognate ligand of TRKB, which increases NMDAR activity and Ca^2+^ influx through NMDAR [59, 60], was also significantly upregulated in the *β-Gal^−/−^* samples (**Figure 4F**). The PI4K antibody binds to isoforms that can be identified by molecular weight. The 230kDa isoform is mostly associated with the ER, while the 55kDa is associated with the PM. Interestingly, only the 55kDa isoform was significantly increased showing that most of this upregulated protein is at the PM [61, 62] (**Figure 4D and S3E-F**).

**Figure 4:**
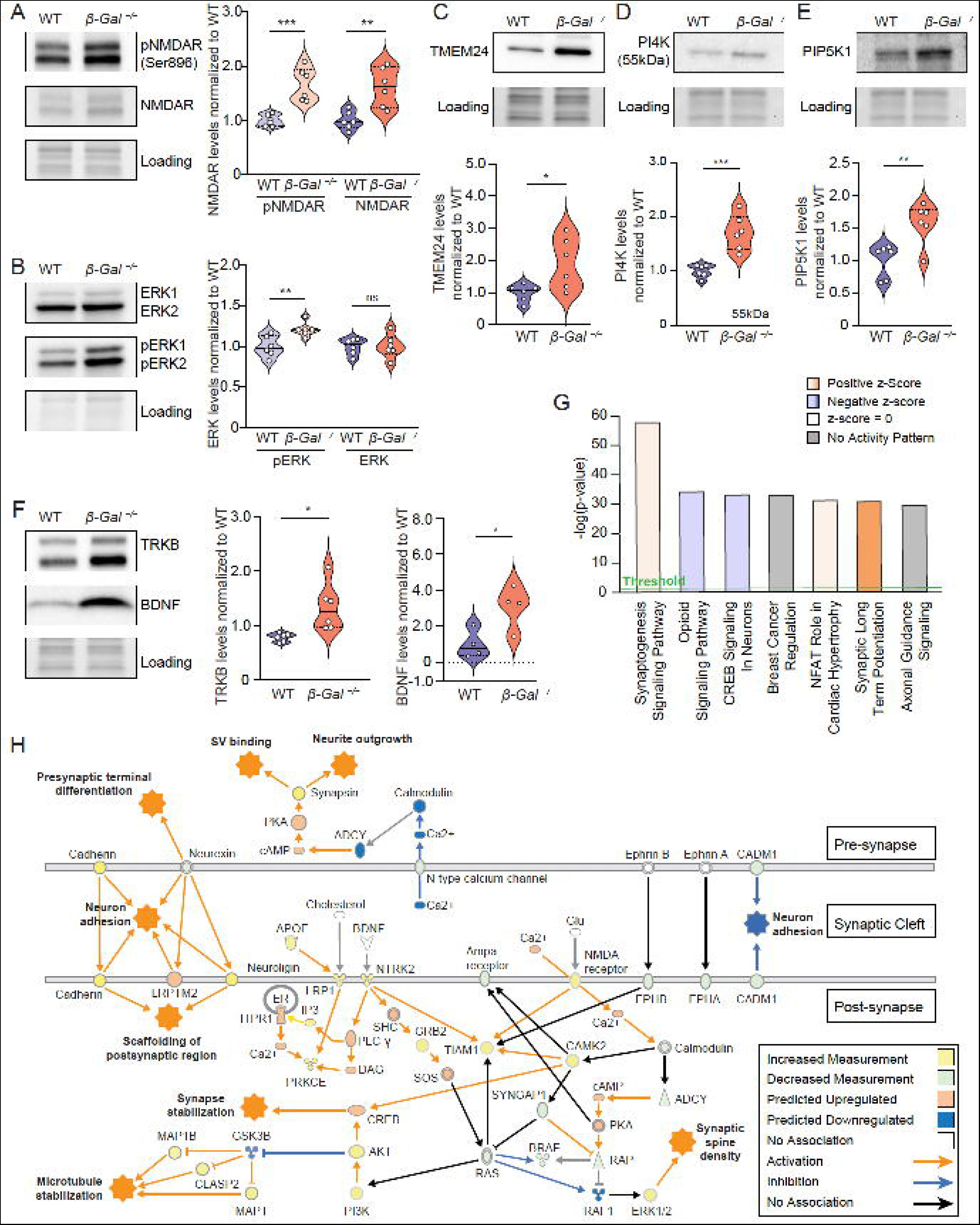
Increased levels of synaptic proteins in *β-Gal^−/−^* ER-PM junctions predict activation of the synaptogenesis-signaling pathway. (A-F) Representative immunoblots and quantification of A) NMDAR-type 1 subunit and pNMDAR (*n* = 6), B) ERK1/2 and pERK1/2 (*n* = 6), C) TMEM24 (*n* = 6), D) PI4K (*n* = 6), E) PIP5K1 (*n* = 6), and F) TRKB (*n* = 6) and BDNF (*n* = 4) in WT and *β-Gal^−/−^* ER-PM junctions isolated from 6-month-old mice. Values are expressed as mean ± SD. Statistical analysis was performed using the Student’s *t*-test; ns: not significant, \**p* < 0.05, \*\**p* <0.01, \*\*\**p* < 0.001. (G) Graphical representation of pathways predicted to be upregulated (orange), downregulated (blue), or dysregulated with no predicted direction (gray), as determined by pathway analysis of proteomics data from ER-PM junctions isolated from 6-month-old WT and *β-Gal^−/−^* mice. Threshold = –log(p=0.05). (H) Network map showing proteins in the synaptogenesis-signaling pathway that were identified in the proteomic analysis of ER-PM junctions from WT and *β-Gal^−/−^* mice.

We next uploaded the proteomics data to the Ingenuity Pathway Analysis (IPA) software to determine cellular processes that are potentially dysregulated in the *β-Gal^−/−^* ER-PM junctions. The top canonical pathways above a threshold of 0.05, according to the Fischer’s Exact test *p*-value, were graphed, and a z-score was used to predict if the pathways were activated (positive z-score, orange) or inhibited (negative z-score, blue), based on the differentially expressed proteins (**Figure 4G**). The pathway most upregulated was the synaptogenesis signaling pathway. Targets for both the presynaptic and postsynaptic terminals were found in the proteomic data, reiterating a role for ER-PM junctions at synapses. In addition, the synaptic LTP pathway was predicted to be highly upregulated (**Figure 4G**). This pathway is mostly associated with increased protein expression at the postsynapse, leading to increased connectivity [22].

We generated a network map of the proteins involved in the synaptogenesis signaling pathway that were predicted in the comparative analysis of ER-PM junctions from WT and *β-Gal^−/−^* mice (**Figure 4H**). The pathways are associated with upregulation of proteins involved in several neuronal processes, including scaffolding of the postsynaptic density, synaptic vesicle binding, neurite outgrowth, and synapse stabilization. These processes are attributed to ERK1/2 activation downstream of NMDAR and TRKB (NTRK2) signaling [57, 63, 64]. We also identified increased Ca^2+^ release from the ER, which concurs with previous findings in our mouse model [42] (**Figure 4H**).Together, these results suggest that GM1 accumulation at *β-Gal^−/−^* ER-PM junctions disturbs synaptic architecture and function, especially at the postsynapse.

### GM1 accumulation in *β-Gal^−/−^* neurons affects the morphology of the dendritic arbor and increases spine formation

To better understand how GM1 accumulation affects the postsynapse, we employed Golgi-Cox staining of the cortex, hippocampus, and cerebellum of 4-month-old WT mice and *β-Gal^−/−^* mice. The number of dendrites in the *β-Gal^−/−^* brain was much less than that in WT brain; this reduction was prominent in the cellular layers of the cortex, the *Cornus Ammonis* of the hippocampus, and the deep cerebellar nucleus (**Figure S4A**).

The cortical neurons of *β-Gal^−/−^* mice contained dendritic structures not found in WT controls, including numerous aberrant neurite outgrowths or ectopic dendrites at the beginning of the dendritic arbor (**Figure S4B**). Another morphologic abnormality was the formation of dendritic beading (**Figure S4C**). This unusual ‘‘string of beads’’ along the dendrites denotes dendritic swelling [65]. The latter often follows dendritic injury and is associated with improper trafficking of molecules between the cell body and the dendrites, leading to a “traffic jam” along the branch and resulting in a bulge [66]. In addition, these neurons had retracted processes and little branching, indicative of an unhealthy neuronal state most likely preceding apoptosis (**Figure S4C**).

Next, we investigated whether GM1 accumulation in the *β-Gal^−/−^* neuronal membranes and ER-PM junctions would affect the number and morphology of dendritic spines. These structures were easily recognized in the pyramidal neurons of the cortex (**Figure S4D**) and hippocampus (**Figure 5A, S4E**). For quantification, we focused on hippocampal pyramidal neurons because that region is involved in LTP, a process that was predicted to be upregulated in *β-Gal^−/−^* ER-PM junctions (**Figure 4G**). The number and density of the spines in the *β-Gal^−/−^* dendritic branches were clearly increased (**Figure 5A, S4E**). For comparative quantification of the number of spines, we used Imaris Imaging software to obtain 3-dimensional (3D) reconstructions of Z-stacked images of Golgi-Cox–stained pyramidal neurons from WT and *β-Gal^−/−^* hippocampi. We traced dendritic branches by using Imaris’ semiautomated filament tracer, which was also used to detect spines along dendritic branches (**Figure 5B**). Quantification of the number of spines within a 10-µm length of dendrite revealed a significant increase in spine density in the *β-Gal^−/−^* pyramidal neurons, compared to that in the WT samples (**Figure 5C**). The number of spines in WT neurons concurred with what has been reported using immunofluorescence imaging [67, 68], thus supporting the validity of this method for spine density quantification. These results suggest that GM1 accumulation at the ER-PM junctions favors spine formation.

**Figure 5:**
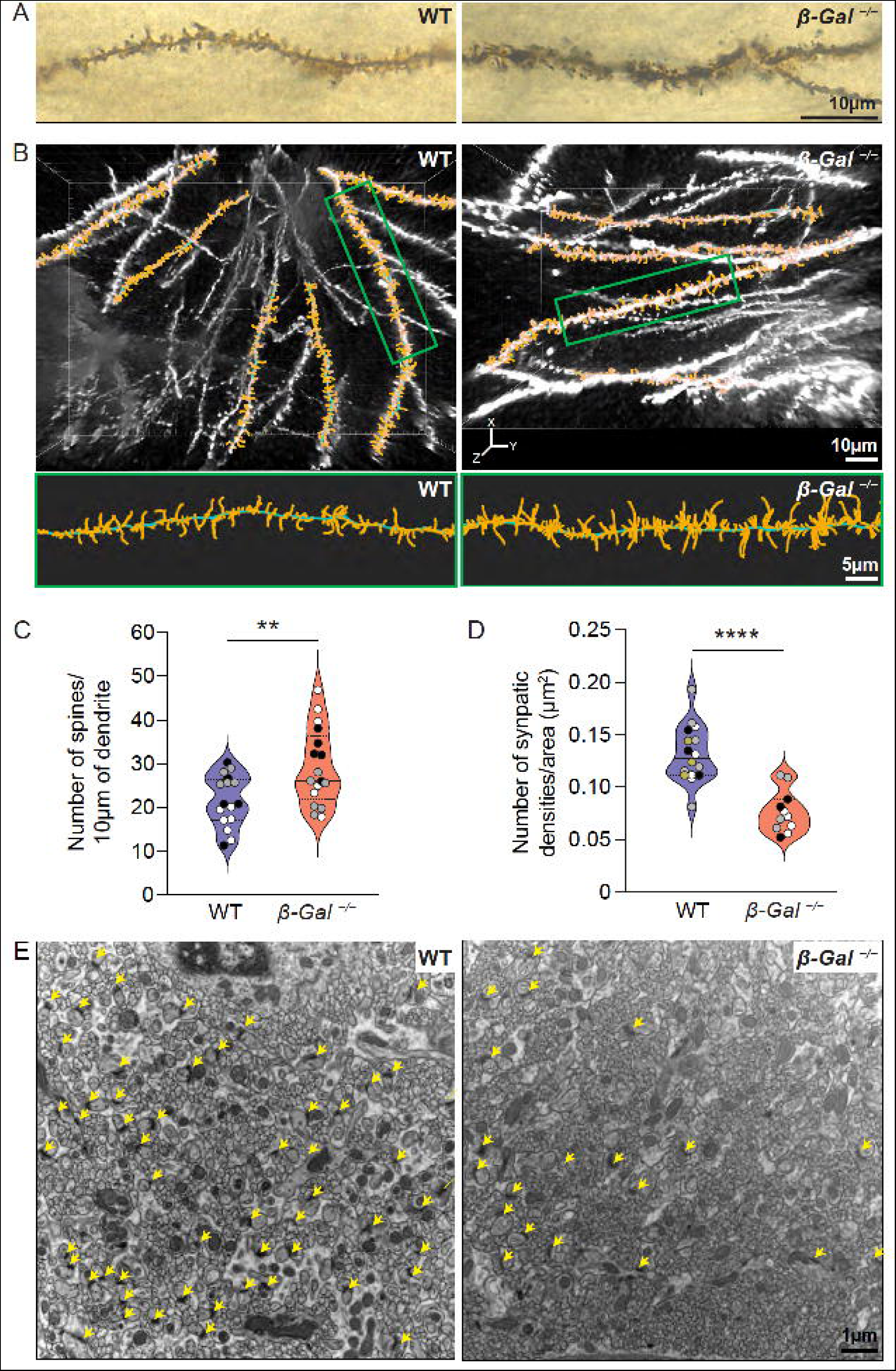
*β-Gal^−/−^* mice have an increased number of dendritic spines but decreased synaptic densities. (A) Golgi-Cox staining of dendritic branches and spines on hippocampal neurons from 6-month-old WT and *β-Gal^−/−^* mice. Scale bar: 10 µm. (B) Three-dimensional (3D) reconstruction of optically sectioned Golgi-Cox–stained dendritic arbors of hippocampal pyramidal cells from 6-month-old WT and *β-Gal^−/−^* mice using Imaris Imaging software. Scale bar: 10 µm. Lower images show 2× zoom of highlighted regions (green boxes) in upper images. Reconstructed dendritic branches (green) and spines (orange) are shown. Scale bar: 5 µm. (C) Quantification of the number of spines per length of dendrite from the 3D reconstruction in B. WT: *n* = 16 and *β-Gal^−/−^*: *n* = 17 neurons; 3 mice, 5-6 images per mouse. Values are expressed as mean ± SD. Statistical analysis was performed using the Student’s *t*-test; \*\**p* < 0.01. (D) Quantification of synaptic densities per field of view of the dendritic arbor in E. Scale bar: 1 µm; n = 16: 4 mice, 3-5 dendritic arbors per mouse (WT), n = 11: 3 mice, 3-5 dendritic arbors per mouse (*β-Gal^−/−^*); each mouse is represented by a different colored filled datapoint (black, grey, white, yellow). Values are expressed as mean ± SD. Statistical analysis was performed using the Student’s *t*-test; \*\*\*\**p* < 0.0001. (E) TEM images of the dendritic arbor of Purkinje cells from 6-month-old WT and *β-Gal^−/−^* mice. Synaptic densities are labeled by yellow arrows.

### Increased number of dendritic spines in the *β-Gal^−/−^* brain does not result in increased synaptic density

An increased number of dendritic spines usually precedes synapse formation [69]. Therefore, we ascertained whether the increased spine density in *β-Gal^−/−^* dendrites would increase the number of synaptic connections. In TEM images, the spines and dendritic branches were readily discernible as cross-sectioned, circular structures within the extensive dendritic arborizations of Purkinje cells (**Figure 5E**). These structures were visibly more numerous in the *β-Gal^−/−^* brain, further confirming the increased formation of spines, albeit their membranes appeared fuzzy and less defined, possibly due to GM1 accumulation (**Figure 5E**). Surprisingly, however, the increased number of spines did not correlate with increased synaptic connections, as determined by the reduced number of electron-dense regions, known as synaptic densities, in the *β-Gal^−/−^* brain (**Figure 5E, yellow arrows, and 5D**).

To confirm that synaptic densities were lost in other regions of the *β-Gal^−/−^* brain, we examined the dendritic arbors of the pyramidal cells in the CA2 region of the hippocampus, where we had quantified spine number by Golgi-Cox staining (**Figure S5A**). In agreement with what we observed in the cerebellum, the CA2 region had more circular processes, indicating an increased spine number, but significantly fewer synaptic densities (**Figure S5A**, **B**).

### GM1 binds pNMDAR and maintains it in an activated state

The observed increase in the levels of total NMDAR-type 1 subunit and pNMDAR in the *β-Gal^−/−^* ER-PM junctions suggested hyperactivation of the receptor at these MCSs. To link these changes in NMDAR levels to the accumulation of GM1 at the ER-PM junctions, we tested whether GM1 interacts with NMDAR-type 1 subunit at these MCSs. Co-immunoprecipitation (co-IP) experiments were performed on the GEM fractions of the ER-PM junctions isolated from WT and *β-Gal^−/−^* brains. Using antibodies against NMDAR-type 1 subunit and pNMDAR, we co-immunoprecipitated GM1, which demonstrated a specific interaction between these molecules in the ER-PM junctions (**Figure S6A, B**).

In the reverse reaction, we used biotinylated cholera toxin B subunit (CTX-B) to pull down GM1 and found that active pNMDAR (**Figure 6A**) preferentially interacted with the ganglioside in the *β-Gal^−/−^* GEMs. In contrast, the non-phosphorylated NMDAR-type 1 subunit appeared to bind GM1 with higher affinity in the WT GEMs (**Figure 6A**). Quantitatively, the ratio of pNMDAR to NMDAR was significantly higher in the GEMs from the *β-Gal^−/−^* ER-PM junctions than in the corresponding WT fractions (**Figure 6B**). The physical interaction between these molecules was further validated by co-staining primary neurons isolated from WT and *β-Gal^−/−^* brains with NMDAR-type 1 subunit antibody and fluorescently labeled CTX-B (**Figure 6C**). NMDAR was increased in *β-Gal^−/−^* neurons, and its distribution pattern overlapped with that of CTX-B, indicating a high degree of co-localization between these molecules (**Figure 6C**). Although the levels of NMDAR and GM1 were quite low in the WT cells, we still detected their co-localization, supporting the notion that these molecules interact under physiological conditions (**Figure 6C**). Together, these results demonstrate that GM1 clusters active pNMDAR at the GEMs, where it may influence the receptor’s synaptically evoked Ca^2+^ signals, leading to increased spine formation.

**Figure 6:**
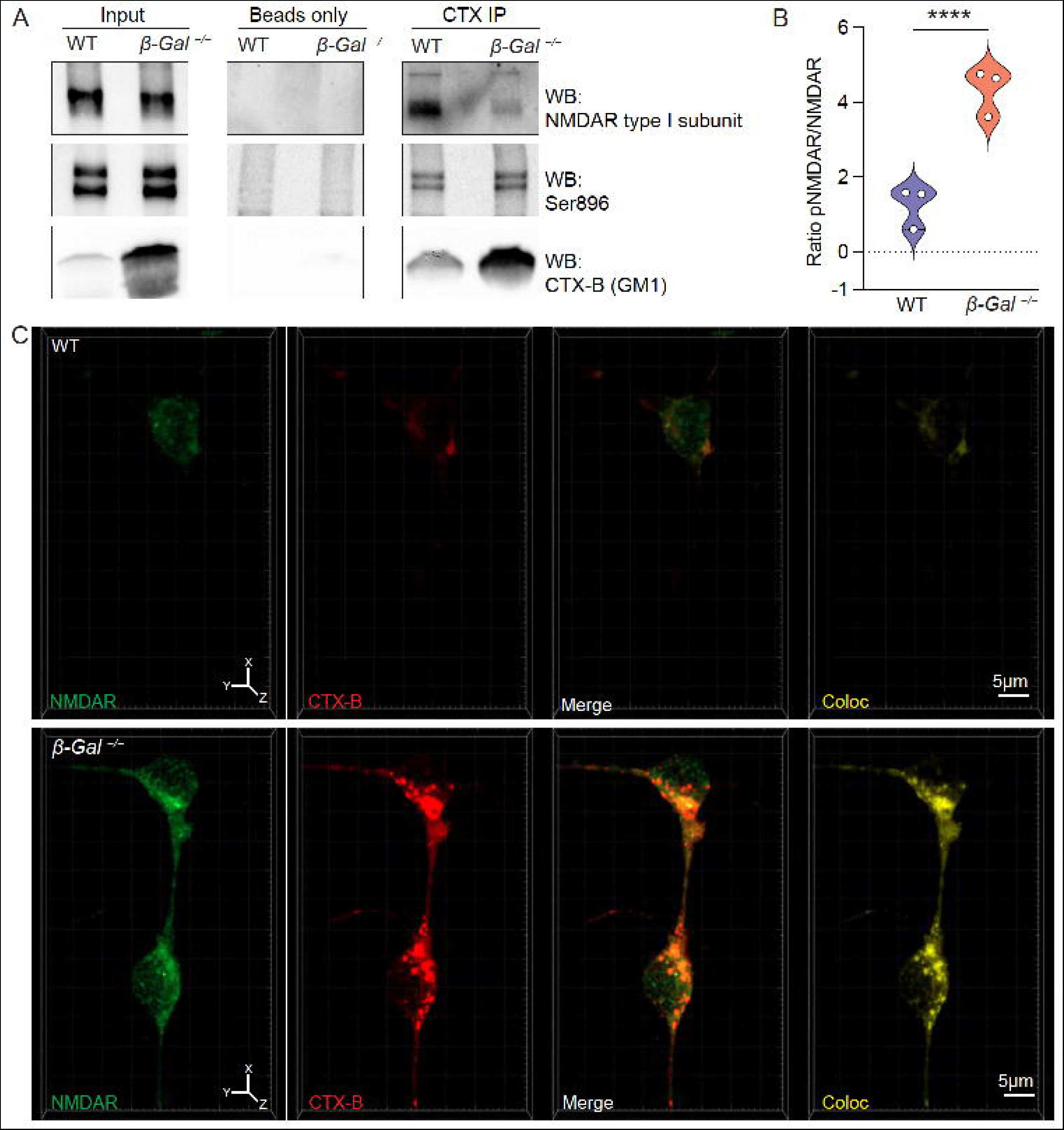
GM1 binds active pNMDAR within *β-Gal^−/−^* ER-PM junctions. (A) Immunoblots of co-IP using cholera toxin B (CTX-B)–conjugated beads to pull down GM1 and bound proteins, NMDAR-type 1 subunit and pNMDAR, in ER-PM junctions isolated from 6-month-old WT and *β-Gal^−/−^* mice. (B) Quantification of the ratio of pNMDAR (Ser896) to NMDAR-type 1 subunit measured in CTX-B co-IP immunoblots; *n* = 3. Values are expressed as mean ± SD. Statistical analysis was performed using the Student’s *t*-test with Welch’s correction; \*\*\*\**p* < 0.0001. (C) Z-stacked immunofluorescence images of primary neurons isolated from P5 WT and *β-Gal^−/−^* brains, stained with CTX-B (GM1; red) and anti-NMDAR-type 1 subunit antibody (green). The colocalization (Coloc) channel shows the generated overlay of GM1 and NMDAR-type 1 subunit staining using Imaris Imaging software. Scale bars: 5 µm.

### GM1 buildup at the ER-PM junctions of *β-Gal^−/−^* neurons affects Ca^2+^ homeostasis and increases the level of calcineurin

The main function of NMDARs is to regulate Ca^2+^ influx into neurons by glutamate stimulation, thereby modulating neuroplasticity [22]. In contrast, GM1 modulates Ca^2+^ flux through its capacity to bind Ca^2+^ and influence the activity of Ca^2+^-binding proteins/receptors [40, 42]. Thus, we hypothesized that GM1 accumulation at the ER-PM junctions and its interaction with pNMDARs at the GEMs provokes a rise in cytosolic Ca^2+^ levels. To test this hypothesis, we measured the cytosolic levels of Ca^2+^ in primary neurons from WT and *β-Gal^−/−^* brains transduced with a lentivirus expressing GCamp6s, a genetically encoded fluorescent Ca^2+^ indicator.

Quantification of the intensity of GCamp6s fluorescence revealed a significantly higher level of Ca^2+^ in *β-Gal^−/−^* primary neurons than in WT controls (**Figure 7A, B**). To link this phenomenon directly to GM1 concentration, we used HeLa cells transduced with GCamp6s that were exogenously loaded with GM1 (**Figure S7A**). GM1-loaded cells showed a significantly higher level of GCamp6s fluorescence than did untreated cells (**Figure S7A, B**).

**Figure 7:**
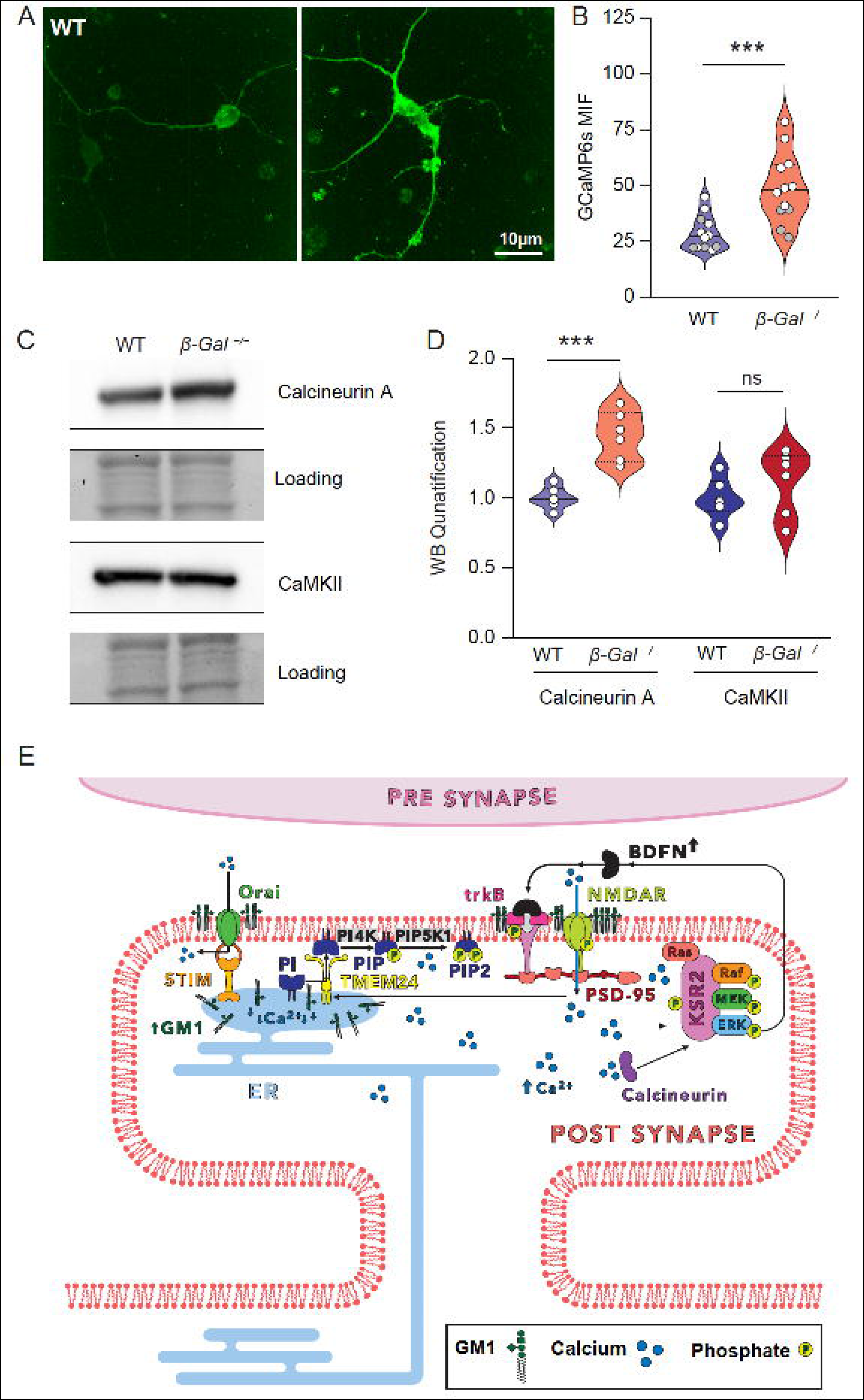
Primary *β-Gal^−/−^* neurons have increased intracellular Ca^2+^ concentration. (A) Fluorescence live-cell Ca^2+^ imaging using the genetically encoded Ca^2+^ sensor, GCamp6s, transduced in primary neurons isolated from the brains of P5 WT and *β-Gal^−/−^* mice. Scale bar: 10 µm. (B) Quantification of the average fluorescence intensity of GCamp6s in primary neurons. *n* = 12, 3 mice with 4 wells per brain. Each mouse is represented by a different colored filled datapoint (black, grey, white). Values are expressed as mean ± SD. Statistical analysis was performed using the Student’s *t*-test; \*\*\**p* < 0.001. (C) Representative immunoblots of calcineurin A and CaMKII in ER-PM junctions isolated from 6-month-old WT and *β-Gal^−/−^* mice. (D) Quantification of immunoblots in C. *n* = 6. Values are expressed as mean ± SD. Statistical analysis was performed using the Student’s *t*-test; ns = not significant; \*\*\**p* < 0.001. (E) Model of the downstream effects of GM1 accumulation in *β-Gal^−/−^* ER-PM junctions: increased cytosolic Ca^2+^ levels through activation of pNMDARs influence the localization of TMEM24 to these MCSs, which mediates the transfer of phosphoinositides (PIs) to the PM. This pool of PIs can then be converted into the signaling molecules, phosphatidylinositol-4-phosphate (PIP) and phosphatidylinositol 4,5-bisphosphate (PIP2), which, in turn, change the synaptic architecture and increase NMDAR PM expression [72, 79]. Concomitantly, increased concentration of cytosolic Ca^2+^ activates calcineurin-dependent ERK signaling, which is necessary for BDNF production. Finally, BDNF activates TrkB, which together with increased NMDARs, induces dendritic spine formation.

Many Ca^2+^-dependent neuronal processes at synapses are mediated by the activity of the kinase CaMKII and the phosphatase calcineurin, both of which are activated after NMDAR-induced Ca^2+^ influx. We found that calcineurin A was significantly upregulated in the *β-Gal^−/−^* ER-PM junctions, but CaMKII was unchanged (**Figure 7C, D**). These results suggest that the increased Ca^2+^ concentration at the ER-PM junctions induces calcineurin-dependent neuronal processes downstream of NMDAR signaling. This phosphatase modulates ERK signaling by interacting with and dephosphorylating the kinase suppressor of Ras 2 (KSR2), thereby influencing KSR2 localization and ERK-scaffold activity in response to Ca^2+^ signals [70] (**Figure 7E**).

## Discussion

In this study, we addressed the role of the sialoglycosphingolipid GM1 in Ca^2+^-mediated processes, where it acts as an integral constituent of the ER-PM junctions. Under neuropathologic conditions that ensue downstream of genetic deficiency of the lysosomal enzyme β-Gal, chronic buildup of GM1 within these MCSs alters the properties of their lipid and protein components, ultimately leading to a synaptopathy associated with abnormal *de novo* spine formation and altered intracellular Ca^2+^ levels. Both these pathogenic events can be attributed to increased levels of NMDAR, TrkB, and BDNF within the *β-Gal^−/−^* ER-PM junctions (**Figure 7E**).

The NMDAR Ca^2+^ channel is highly expressed in excitatory postsynapses, where it promotes LTP and spine formation, especially in pyramidal cells of the hippocampus [22, 71]. In addition, NMDAR-induced Ca^2+^ influx into the cytosol of neurons influences the localization of the lipid transporter TMEM24 to the ER-PM junctions, although its function as a PI transporter has not yet been demonstrated in neurons [34]. It is possible that TMEM24 relocates to the *β-Gal^−/−^* ER-PM junctions, facilitating the transport of PIs to these MCSs during synaptogenesis and spine formation, however this hypothesis needs to be further investigated. PI4K and PIP5K1 then convent the pool of PIs within the MCSs into PIP and PIP2, both of which are potent neuronal-signaling molecules that modulate the activity and PM expression of NMDARs at the synapse [72] (**Figure 7E**). One important function of GM1 is interacting with phosphorylated membrane receptors, thereby prolonging their activation. We found that GM1 interacts specifically with pNMDARs at Ser896, a site that increases the receptor’s expression at the PM and holds this Ca^2+^ channel in an active conformation at the ER-PM junctions. This interaction perpetuates a vicious cycle of abnormal Ca^2+^ influx, which activates ERK signaling, leading to excessive spine formation without increasing connectivity to the presynaptic axons (**Figure 7E**). These results parallel what we have previously shown in the MAMs, where GM1 binds IP3R-1 in an active phosphorylated state, increasing ER Ca^2+^ efflux through the IP3R-1/VDAC1/GRP75 Ca^2+^ megapore [42]. In addition, binding GM1 to phosphorylated, active TrkB at the PM favors its interaction with BDNF, which in turn induces neuritogenesis downstream of ERK signaling [47]. BDNF also matures synaptic spines in the hippocampus and increases their numbers [63]; both of these processes may potentiate the effects of NMDARs.

We found that the pNMDAR-mediated increase in intracellular Ca^2+^ levels activated calcineurin rather than CaMKII. In neurons, calcineurin dampens intracellular Ca^2+^ levels by dephosphorylating and inactivating Ca^2+^ channels at the PM [73], which eventually decreases synaptic spine density [74, 75]. Calcineurin also modulates ERK signaling at the PM by interacting with and dephosphorylating the KSR2, thereby influencing KSR2 localization and ERK-scaffold activity in response to Ca^2+^ signals. This promotes BDNF production and ultimately contributes to the increased number of spines [70]. In our *β-Gal^−/−^* model, a potential explanation for the perceived opposite role of calcineurin in spine formation is that high levels of the phosphatase at the ER-PM junctions counteract the increase in the number of non-connecting spines. However, chronic activation of pNMDARs in complex with GM1 at ER-PM junctions outcompetes calcineurin’s capacity to limit the number of spines.

Altered numbers of synaptic spines has been reported in several neurologic conditions, including common neurodegenerative diseases, autism spectrum disorders, and Fragile X syndrome [76–78]. In Fragile X syndrome, the combined deregulation of metabotropic glutamate receptors and increased sensitivity to BDNF signaling contribute to excessive dendritic branching and spine formation [76, 77]. These findings further support the idea that increased NMDARs and BDNF most likely cause excessive spine formation in *β-Gal^−/−^* mice.

A common element that adds to the complexity of MCS-regulated pathways is altered Ca^2+^ signaling caused by GM1 accumulation at both the MAMs and ER-PM junctions, two MCSs with similar structural and functional characteristics. Interestingly, we observed an increased number of triple-contact sites involving the ER, PM, and mitochondria in *β-Gal^−/−^* neurons compared to WT neurons. These sites may bridge the activity of the MAMs and ER-PM junctions, potentiating the effects of increased Ca^2+^ concentration at the synapses of the *β-Gal^−/−^* mice.

These studies put forward previously unknown functions of GM1 within neuronal ER-PM junctions in the regulation of membrane dynamics at the synapses. Given that β-Gal-mediated catabolism of GM1 occurs in lysosomes and that these organelles tether/fuse with virtually all other intracellular membranes and the PM, understanding the effects of GM1 accumulation at MCSs involving the lysosomal membrane warrants further investigation.

## Acknowledgments

We thank Dr. Xiaohui Qiu for help with the production of lentiviruses. A.d’A. holds the Jewelers for Children Endowed Chair in Genetics and Gene Therapy. This work is supported by the NIH grants DK052025, CA021764, and grants from the Assisi Foundation of Memphis, the National Tay-Sachs and Allied Diseases (NTSAD), and the American Lebanese Syrian Associated Charities (ALSAC). The content is solely the responsibility of the authors and does not necessarily represent the official views of the National Institutes of Health.

## Author Contributions

A.d.A. conceived and designed the study and edited the manuscript; J.A.W. designed the experiments, collected and analyzed data, and drafted the manuscript; I.A. coordinated the experimental design and edited the first draft; D.v.d.V. edited the Star Methods and formatted the figures; C.G.R. performed TEM analyses and assisted in data interpretation; Y.C. produced the lentivirus constructs; A.M. performed proteomic analyses; L.E.F. reviewed the manuscript and assisted with experiments; H.H. performed histologic analyses; and E.G. helped collect animal tissues and maintained the animal colony. All authors have read and agreed to the published version of the manuscript.

## Declaration of interests

The authors declare no competing interests.

## STAR METHODS

### RESOURCE AVAILABILITY

#### Lead contact

Further information and requests for resources and reagents should be directed to and will be fulfilled by the lead contact, Alessandra d’Azzo.

#### Materials availability

Plasmids used in this study are readily available at Addgene. This study did not generate new unique reagents.

#### Data and code availability

Proteomics data have been deposited to the ProteomeXchange Consortium *via* the PRIDE partner repository with the dataset identifier (PXD042994).

### EXPERIMENTAL MODEL DETAILS

#### Animal Model

Animals were housed in a fully AAALAC (Assessment and Accreditation of Laboratory Animal Care)-accredited animal facility with controlled temperature (22°C), humidity, and lighting (alternating 12–h light/dark cycles). Food and water were provided *ad libitum*. All procedures in mice were performed according to animal protocols approved by the St. Jude Children’s Research Hospital Institutional Animal Care and Use Committee (IACUC) and NIH guidelines. The *β-Gal^−/−^* murine model was generated as previously described [52]. WT and *β-Gal^−/−^* mice (C57BL/6J background), aged 5 days to 6 months, were used for experiments and are described in the METHODS DETAILS. Both male and female mice were used in all experiments without any apparent sex differences.

#### Cell lines

##### HeLa cells and HEK293T cells

were maintained in culture in Dulbecco’s Modified Eagles Medium (DMEM, Gibco) supplemented with penicillin/streptomycin (100 μg/mL) (Gibco), 2 mM GlutaMAX (Gibco), and 10% fetal bovine serum (FBS; Thermo Fisher) and incubated at 37°C in 5% CO_2_.

##### Primary neurons

were isolated from WT and *β-Gal^−/−^* mice (as described in METHODS DETAILS), plated on poly-D-lysine–coated glass coverslips, cultured in MACS® Neuro Medium (Miltenyi) containing MACS® NeuroBrew®-21 (Miltenyi), and incubated at 37°C in 5% CO_2_.

### METHODS DETAILS

#### Transmission electron microscopy analysis

Mouse brains were isolated and fixed in 0.1M cacodylate buffer containing 2.5% glutaraldehyde and 2% paraformaldehyde (PFA). Fixed brains were embedded in 4% low–melting point agarose (Sigma Aldrich) and cut into 100-µm sagittal sections on a VT1000 S vibratome (Leica). Samples were postfixed in reduced osmium tetroxide and contrasted with aqueous uranyl acetate. Dehydration was performed by a series of ethanol baths, ascending in concentration from 70% to 100%, followed by 100% propylene oxide. Samples were infiltrated with EmBed-812 and polymerized at 60°C. Embedded samples were sectioned at ~70 nm on a Leica (Wetzlar) UC-7 ultramicrotome and examined in a ThermoFisher Scientific (Hillsboro) TF20 transmission electron microscope at 80 kV. Digital micrographs were captured with an Advanced Microscopy Techniques (AMT; Woburn) imaging system. Whole-cell montages were acquired with the same AMT system controlled by serialEM software (Mastronarde, 2005). Montages were stitched and exported as .tif files by using IMOD software (**Kremer et al. 1996**). Unless otherwise indicated, all reagents were from Electron Microscopy Sciences (Hatfield). For ER-PM junction quantification, Purkinje cell bodies were imaged by montage around the entire soma. The total length of the PM and the fragments of ER within 35 nm of the PM were traced and measured. To quantify synaptic density, 3 consecutive images were taken of the dendritic arbor of Purkinje cells, and the average numbers of synaptic densities per area were measured. In the hippocampus, consecutive images were taken of the dendritic arbors parallel to the pyramidal cell layer in the CA2 region to determine the average number of synaptic densities per area.

#### Production of lentivirus vector

Lentiviral vector preparations were produced by co-transfection of HEK293T cells with a mixture of 4 plasmids (13.5 µg pCAG-kGP1-1R, 4.5 µg pCAG4-RTR2 [80], 4.5 µg envelope plasmid pCAG-VSVG, and 22.5 µg vector plasmid pCL20-GFP-MAPPER or pGP-CMV-GCaMP6s) using the calcium phosphate precipitation method: 9 × 10^6^ HEK293T cells were seeded onto 15-cm tissue culture dishes 24 h before transfection. The mixture of plasmids was diluted to a total volume of 450 µL with ddH_2_O. Then 50 µL 2.5M CaCl_2_ was added, and the mixture was incubated for 5 min at room temperature (RT); 2× Hank’s balanced salt solution (HBSS; 500 µL) (50mM HEPES, pH 7.05; 280mM NaCl, 1.5mM Na_2_PO_4_) was added dropwise while vortexing, and the mixture was incubated for 10 min at RT. The final mixture (1 mL DNA-CaPO_4_-HBSS) was added dropwise to a 15-cm dish containing 25 mL DMEM supplemented with penicillin/streptomycin (100 μg/mL), 2 mM GlutaMAX, and 10% FBS and then mixed well by swirling/tilting the dish.

The cells were incubated at 37°C in 5% CO_2_ for 18 h. The medium was exchanged for Neurobasal medium [-] L-glutamine (Gibco, Cat. No. 21103-049) without serum, and the cells were incubated for 24 h. The next day, the supernatant containing virus was collected, cleared by 500 ×*g* for 5 min at RT and filtered through a 0.45-µm filter. The virus was concentrated (~100×) by ultracentrifugation at 112,400 ×*g* at 4°C for 90 min, and the viral pellet was resuspended in Neurobasal medium without serum overnight at 4°C. Aliquots were snap-frozen on dry ice and stored at –80°C.

#### Isolation of neurons

P5 WT mice and *β-Gal^−/−^* mice were used to set up neuronal cultures. The pups were euthanized by CO_2_, and their brains were collected in dissection media (500 mL HBSS, 5 mL 1M HEPES, 6.67 mL 45% glucose, penicillin (100 U/mL), and streptomycin [100 µg/mL]). The cerebellum, olfactory bulbs, and meninges were removed and discarded. The remaining brain dissociated using the Neural Tissue Dissociation Kit – Postnatal Neurons (Miltenyi Biotec). Brains were placed in a gentleMACS^TM^ C tube (Miltenyi Biotec) with the supplied enzymes and dissociated on a gentleMACS^TM^ Octo Dissociator with Heaters (Miltenyi Biotec) on the “NTDK_37C” program. The brain suspension was passed through a 1000-μL pipette tip 5× with 1mL of 5% bovine serum albumin (BSA)/PBS to fully homogenize it and filtered on a 70μm cell strainer. Tubes were centrifuged at 300 ×*g* for 5 min, and the supernatant was removed. The pellet were resuspended in 80 μL cold 5% BSA/PBS and 20 μL non-neuronal cell biotin antibody (Miltenyi Biotec; Neuron Isolation kit) and incubated at 4°C for 5 min. The pellet was resuspended in 800 μL cold 5% BSA passed through an LS column (Miltenyi Biotec) on a magnetic strip to remove non-neuronal cells. The flow through, which contained the neurons, was collected and centrifuged at 300 ×g for 5 min at 4°C. Cells were plated in poly-D-lysine–coated 2-well glass coverslip chambers (LabTak) using neuronal basal media containing B-27 supplements and incubated at 37°C in 5% CO_2_.

#### MAPPER analysis *in vitro*

HeLa cells and primary neurons were maintained in culture in 2-well glass bottom slides (LabTek). The cells were grown until confluency and transduced with 10 µL MAPPER lentivirus in 500 µL media for 4 h at 37°C in 5% CO_2_. The cells were washed 2× with PBS, fresh media was added, and the slides were incubated overnight at 37°C in 5% CO_2_. For HeLa cells, 1 µg/µL GM1 (Advanti) was added overnight in 0.5% serum media to load cells with GM1. Live-cell confocal imaging was performed using the Marianas 2Axio Observer microscope (Zeiss) at 37°C/5% CO_2_ for 10 min in the green fluorescence channel (488 nm), capturing images every 15 s. After 1 min of imaging, 1µM thapsigargin was added to induce ER-PM junction formation, or 1µM DMSO (control) was added to the wells, and imaging was continued for 9 min.

#### Isolation of ER-PM junctions

Mice were euthanized by CO_2_ and brains were isolated. Brain hemispheres were separated on ice. Each hemisphere was homogenized in a 2-mL Kontes Dounce Homogenizer (Kimble) in 1 mL homogenizer solution (0.32M sucrose, 1mM NaHCO_3_, 1mM MgCl_2_, 0.5mM CaCl_2_, protease and phosphatase inhibitors) with 15 strokes of a loose-fitting glass pestle (Kimble).

##### Nuclear removal steps

Homogenates were transferred to a 15-mL Falcon tube and diluted up to 10% w/v (e.g., 0.225 g = 2.25 mL) with homogenizer solution and centrifuged at 1400 ×*g* for 10 min at 4°C. Supernatant was transferred to a 30-mL glass Corex tube on ice, and pellets were resuspended in the same 10% w/v homogenizer solution and homogenized in a 2-mL Kontes Dounce Homogenizer with 6 strokes of a tight-fitting glass pestle (Kimble). Homogenates were transferred to a 15-mL Falcon tube and centrifuged at 710 ×*g* for 10 min at 4°C. The resulting pellet containing nuclear and cellular debris was discarded.

##### ER-PM-Mito enrichment steps

Supernatants were combined in a 30-mL glass Corex tube after the previous nuclear removal steps and centrifuged in a Sorvall centrifuge at 13,800 ×*g* for 10 min at 4°C. Pellets were resuspended in the same 10% w/v homogenizer solution and homogenized in a 2-mL Kontes Dounce Homogenizer with 6 strokes of a tight-fitting glass pestle. Homogenates were transferred to a 30-mL glass Corex tube and centrifuged in a Sorvall centrifuge at 13,800 ×*g* for 10 min at 4°C. Homogenization and centrifugation steps were repeated, and pellets (enriched in PM, bound ER, and mitochondria [Mito]) were collected for the sequential steps. After the 3 centrifugations, supernatants were combined in a 13-mL Ultra-Clear centrifuge tube (Beckman) and ultracentrifuged in a Beckman Ultracentrifuge at 100,000 ×g for 1 h at 4°C. The resulting supernatant contained the cytosolic fraction, and the pellet contained the unbound ER fraction. **Figure S2A** shows representative immunoblots of the fractions collected during the nuclear removal and ER-PM-Mito–enrichment steps, probed with the appropriate organelle markers: laminin A/C (nucleus), calnexin (ER), N-cadherin (PM), lactose dehydrogenase (LDH; cytosol), and TOMM20 (mitochondria).

##### Sucrose gradient

The pellets (enriched in PM, bound ER, and mitochondria) were homogenized in 2 mL 11% w/v sucrose in ER-PM buffer (5 mM Bis-Tris, 0.5 mM NaHCO_3_, 0.2 mM EDTA, protease inhibitors, pH 6) in a 2-mL Kontes Dounce Homogenizer with 10 strokes of a tight-fitting glass pestle. Homogenate was layered on a discontinuous sucrose gradient containing 2 mL 53% w/v, 43% w/v, and 4 mL 38% w/v sucrose in ER-PM buffer in a 13-mL Ultra-Clear centrifuge tube and centrifuged at 100,000 ×*g* for 2.5 h at 4°C. Centrifugation resulted in 3 distinct bands: ER-PM junctions (top), mitochondria (middle), and PM (bottom). The top band was collected and resuspended in 7 mL ER-PM wash buffer (225mM mannitol, 75mM sucrose, 0.5mM EGTA, 30mM Tris-HCl, pH 7.4) and centrifuged at 13,000 ×*g* for 10 min at 4°C. Supernatant was transferred to a 13-mL Ultra-Clear centrifuge tube and centrifuged at 100,000 ×*g* for 40 min at 4°C. The resulting pellet contained the ER-PM junctions. The bottom band (PM) was collected and resuspended in 15 mL ER-PM wash buffer and centrifuged at 13,000 ×*g* for 10 min at 4°C. Supernatant was transferred to a 32-mL Ultra-Clear centrifuge tube (Beckman) and centrifuged at 48,000 ×g for 20 min at 4°C. The resulting pellet contained the PM fraction. Pellets containing ER, PM, and ER-PM junctions can be resuspended in RIPA buffer (50mM Tris-HCl, pH 7.5; 150-mM NaCl, 0.1% SDS, 1% DOC, 1% TritonX-100, protease and phosphatase inhibitors) for Western blot and proteomic analyses, in HPLC-grade water for HP-TLC analysis or separated further into the GEM fraction (see below). The bands corresponding to the ER-PM junctions and PM fractions (**Figure S2B**) were individually collected and further centrifuged; the resulting pellets were then used for further experimentation.

#### GEM isolation of ER-PM junctions

Pellets containing ER, PM, and ER-PM junctions were resuspended in 600 µL extraction buffer (25mM HEPES, pH 7.5; 0.15M NaCl, 1% TritonX-100, and protease and phosphatase inhibitors) and incubated on ice for 20 min. Lysates were passed through a 22-G syringe needle 15× and then centrifuged at 14,000 ×g for 2 min at 4°C. Supernatants were collected and re-centrifuged for 2 min to remove any remaining soluble material. Pellets were then combined and solubilized in solubilizing buffer (50mM Tris-HCl, pH 8.8; 5mM EDTA, 1% SDS).

#### Spectral-count proteomic analysis

ER-PM junction fractions were resuspended in Lameli buffer (BioRad) and run on Mini-PROTEAN 4%-20% TGX gels (BioRad) until the sample entered the gel. Bands were then excised. The proteins in the gel bands were reduced with DTT to break disulfide bonds, and the Cys residues were alkylated by iodoacetamide to allow the recovery of Cys-containing peptides. The gel bands were then washed, dried in a speed vacuum, and rehydrated with a buffer containing a protease to enable the protease to enter the gel. Typically, trypsin enzyme is used for digesting proteins, but additional enzymes may be used to achieve better coverage of protein sequence. In this experiment, each band was digested independently with trypsin and analyzed by mass spectrometry (MS). Raw data files for each sample were combined prior to database search. The peptide samples were loaded on a nanoscale capillary reverse-phase C18 column (2.7 µm C18, 100 mm) by an HPLC system (Waters ACQUITY UPLC or Thermo EASY-nLC 1000) and eluted by a gradient (~60 min). The eluted peptides were ionized and detected by an inline mass spectrometer (Thermo Fusion). MS data were collected first (in ~0.5 s), followed by sequential isolation for MS/MS analysis (each in ~0.1 s, totaling ~2 s) acquisition of the top-10 abundant ions and additional target Mz representing user-selected sites. This process (~2.5 s) was repeated over the entire elution chromatography gradient, acquiring more than 75,000 MS/MS spectra during a 180-min elution.

Database searches were performed using Sequest search engine in Thermo Proteome Discoverer 1.4 (v1.4.0.288) and PEAKS Studio 8.5 (Build 20180507) software packages. All matched MS/MS data were filtered by mass accuracy and matching scores to reduce the protein false-discovery rate to ~5%. Modification sites were determined by dynamically assigning related mass addition to all possible amino acid residues during the database search. Peptide assignments were further analyzed by Ascore, pRS score, and de novo sequencing programs and were subjected to manual examination. All proteins identified in a gel lane were combined. The total number of spectra, namely spectral counts, matching individual proteins may reflect their relative abundance in one sample, after the protein size is normalized. Moreover, the spectral count is useful for comparing the levels of the same protein across multiple samples (e.g., control and immunoprecipitated samples). Modified residues were further validated by an independent de novo sequencing of raw spectra and confirmed based on the unambiguous assignment of characteristic site-specific fragment ions. In addition, the ion intensities of modified peptides and corresponding unmodified counterparts were compared to evaluate the stoichiometry of the modification events.

#### Golgi-Cox immunohistochemical staining

Golgi-Cox staining was performed using FD Rapid GolgiStain Kit (FD NeuroTechnologies). In short, mice were euthanized by CO_2_ and brains were quickly isolated. The brains were rinsed with Milli-Q water to remove blood and immersed in impregnation solution (FD NeuroTechnologies) for 2 weeks at RT in the dark. The brains were transferred to development solution (FD NeuroTechnologies) for 72 h at RT in the dark. Brains were then flash-frozen in liquid nitrogen and cut into 100- to 200-µm sections on a cryostat. The brain sections were mounted on gelatin-coated slides and developed with staining solution (FD NeuroTechnologies). Sections were then dehydrated in 50%, 75%, and 95% ethanol for 4 min each. The sections were further dehydrated in 100% ethanol 4× for 4 min each. Sections were then cleared with xylene 3× for 4 min each and mounted in Cytoseal (Fischer) with a coverslip.

#### 3D reconstruction of dendritic spines

For 3D reconstruction of Golgi-Cox–stained hippocampal pyramidal cells, Z-step brightfield images (60× oil immersion lens) were taken using the Lionheart FX automated microscope (Biotek), which enabled us to visualize dendritic spines. We performed linear deconvolution using ImageJ software and exported images to Imaris Imaging software for 3D reconstruction of the cells/tissue. Dendritic branches and spines were traced using the Filament Tracer software in a semiautomated fashion. Similar thresholds were used for spine-density seeding to enable the software to reconstruct the spines. Data readouts show the number of spines per 10-µm length of the dendritic branch.

#### GM1 isolation and quantification by HPTLC

To extract the lipids from the ER-PM junction fractions, we resuspended the ER-PM pellets in 3× volume of HPLC-grade water in an Eppendorf tube. An 8× volume of methanol was added to each sample (8:3 volume CH_3_OH:dH_2_O) and vortexed at RT. Chloroform equaling half the volume used for methanol was added to each sample (4:8:3 volume CHCl_3_:CH_3_OH:dH_2_O) and vortexed at RT. The mixture was then centrifuged at 1200 ×*g* for 15 min at RT. The volume of supernatant was measured and transferred into a fresh Eppendorf tube. Water (0.173× the volume) was added to the supernatant, vortexed, and centrifuged at 1200 ×*g* for 15 min at RT. The upper polar phase was collected in a new tube and evaporated at 48°C overnight, until the lipid pellet remained. To analyze the GM1 content, we resuspended lipid pellets in 20-50 µL chloroform solution (4:8:3, CHCl_3_:CH_3_OH:dH_2_O). A sample (5 µL) was loaded onto a 10 × 20 cm silicone-coated TLC plate (Millipore) using the Automatic TLC Sampler ATS 4 (Camag). GM1 (1 µg/µL) isolated from bovine brain (Sigma, Cat no. G7641) in the 4:8:3, CHCl_3_:CH_3_OH:dH_2_O solution was plated as a standard. Plates were developed in an Automatic Developing Chamber ADC2 (Camag) using 60:35:8, CHCl_3_:CH_3_OH:0.25% KCl as development solution. Plates were then sprayed with resolving solution (2% resorcinol, 80% HCl, 5% 0.1M copper sulfate) and heated on a hot plate for 30 min at 95°C, until bands appeared. Plates were imaged on the TLC Visualizer 2 (Camag), and bands containing GM1 were quantified using the GM1 standard.

#### Immunoblotting and immunoprecipitation

For immunoblotting, samples were homogenized with RIPA buffer, and protein concentrations were determined using the Pierce BCA kit (Thermo Scientific). Proteins were separated by SDS– PAGE on precast Mini-PROTEAN 4%-20% TGX gels (BioRad), under reducing conditions. They were then transferred to a polyvinylidene difluoride (PVDF) membrane (Millipore). Membranes were incubated for 1 h in blocking buffer (5% dry milk in TBS-Tween) at RT and subsequently probed with the specific primary antibody diluted in 5% BSA/TBS-Tween overnight at 4°C at the following concentrations: anti-BDNF, 1:1000 (Abcam); anti-pan calcineurin A, 1:1000 (Cell Signaling); anti-calnexin, 1:1000 (Novus Biologicals); anti-CaMKII, 1:500 (Santa Cruz); anti-ERK1/2, 1:1000 (Cell Signaling); anti-GM1, 1:1000 (Abcam); anti-laminin A/C, 1:1000 (Cell Signaling); anti-LDH, 1:1000 (Chemicon); anti-N-cadherin, 1:1000 (BD Biosciences); anti-NMDAR-type 1 subunit, 1:1000 (Cell Signaling); anti-ORAI1, 1:500 (Abcam); anti-pERK1/2, 1:1000 (Cell Signaling); anti-PI4K, 1:1000 (Cell Signaling); anti-PIP5K1A, 1:1000 (Cell Signaling); anti-pNMDAR (Ser896), 1:1000 (Invitrogen); anti-STIM1, 1:1000 (Cell Signaling); anti-STIM2, 1:1000 (Cell Signaling); anti-TMEM24, 1:1000 (Fitzgerald); anti-TOMM20, 1:1000 (Santa Cruz); anti-TRKB, 1:1000 (Cell Signaling); anti-VAPA, 1:1000 (Bethyl); anti-VAPB, 1:1000 (Bethyl); and CTX-B HRP-conjugated, 1:5000 (Invitrogen). Blots were then washed 3× with TBS-Tween for 5 min and incubated with the appropriate secondary antibody for 1 h at RT. Blots were washed 3× with TBS-Tween for 5 min and developed using the Clarity Max^TM^ Western ECL Substrate (BioRad).

For immunoprecipitation, the GEMs isolated from ER-PM junctions were diluted in 500 µL RIPA buffer containing 2× protease inhibitors. Lysates were precleared with 30 µL Pansorbin cells (Millipore Sigma) for 1 h at 4°C. Lysates were washed with RIPA and incubated with anti-NMDAR-type 1 antibody (1:50), anti-pNMDAR (Ser 896) (1:50), or CTX-B biotin-conjugated (1:100). The samples were then incubated overnight at 4°C under agitation. 20 µL Protein A beads (Cell Signaling) or 5 µL Streptavidin Magnetic beads (Vector Lab) for CTX_B were incubated for 1 h at 4°C. Samples were centrifuged at 10,000 ×*g* for 30 s at 4°C and placed on a magnetic strip, and supernatant was removed. Pellets were washed 3× with RIPA buffer and resuspended in Laemmli sample buffer (BioRad) before immunoblotting on precast Mini-PROTEAN 4%-20% TGX gels. Proteins were then transferred onto PVDF membranes and probed with HRP–conjugated CTX-B to visualize GM1.

#### Immunofluorescent cell imaging

Primary Neurons were isolated from P5 WT mice and *β-Gal^−/−^* mice were cultured for 10 days and permeabilized and fixed using cold acetone for 5 min at −20°C. Cells were washed 3× with PBS and blocked in 0.25% Casine in PBS for 1 h at RT. Cells were incubated in 1:100 anti-NMDAR-type 1 subunit antibody and 1:1000 CTX-B Alexa Fluor^TM^ 555-conjugated overnight at 4°C. Cells were washed 3× in PBS and incubated with donkey anti-Rabbit Alexa Fluor^TM^ 488-conjugated IgG secondary antibody (Invitrogen) for 1 h at RT. Cells were washed and mounted on a coverslip using ProLong™ Gold Antifade Mountant with DNA Stain DAPI (ThermoFisher). The amount of colocalized area was calculated using Imaris.

#### Ca^2+^ imaging in vitro

HeLa cells and primary neurons were put into culture in a 2-well glass bottom slide (LabTek) using the methods described above and 1 mL of the appropriate media described. Media was aspirated, and cells were transduced with 10 µL Lenti-Gcamp6s construct in 500 µL media for 4 h at 37°C in 5% CO_2_. Media was removed and cells were washed 2× with PBS for 5 min. Media (1 mL) was added to the cells, and slides were incubated at 37°C in 5% CO_2_ overnight. To load HeLa cells with GM1, 1 µg/µL GM1 was added overnight in 0.5% serum media. Slides were then imaged on the Marianas 2Axio Observer microscope at 37°C in 5% CO_2_ in the green fluorescence channel (488 nm). Z-stack imaging was performed on the entire cell body, and maximum-intensity projections were performed on 2D flattened images.

### QUANTIFICATION AND STATISTICAL ANALYSES

Statistical analyses were performed using GraphPad Prism. Quantitative data are presented as mean ± SD, or as mean ± 95% confidence interval (MAPPER experiment). For comparisons between 2 groups, Student’s *t*-tests (paired or unpaired, 2-tailed) were performed. For small sample sizes (n <6), a Welch’s correction was applied to the Student’s *t*-test. For comparisons of 3 or more groups, one-way or two-way ANOVAs were performed. P-values <0.5 were considered statistically significant. Numbers of samples are indicated in the figure legends.

### KEY RESOURCES TABLE

| REAGENT or RESOURCE | SOURCE | IDENTIFIER |
| --- | --- | --- |
| Antibodies |  |  |
| Rabbit monoclonal anti-BDNF antibody | Abcam | ab108319 |
| Rabbit anti-pan calcineurin A antibody | Cell Signaling | 2614S |
| Rabbit polyclonal anti-calnexin antibody | Novus Biologicals | NB100-1965 |
| Mouse monoclonal anti-CaMKII alpha (A-1) antibody | Santa Cruz | Sc-13141 |
| Rabbit monoclonal anti-p44/42 (ERK1/2) antibody | Cell Signaling | 4695S |
| Rabbit polyclonal anti-Ganglioside GM1 antibody | Abcam | ab23943 |
| Rabbit anti-Laminin A/C antibody | Cell Signaling | 2032S |
| Goat anti-LDH antibody | Chemicon Intl. | AB1222 |
| Mouse anti-N-Cadherin antibody | BD Biosciences | 610921 |
| Rabbit monoclonal anti-NMDA Receptor 1 antibody | Cell Signaling | 5704S |
| Rabbit polyclonal anti-ORAI1 antibody | Abcam | ab86748 |
| Mouse monoclonal anti-phospho-p44/42 (ERK1/2) (thr202/Tyr204) antibody | Cell Signaling | 9106S |
| Rabbit anti-PI4 Kinase antibody | Cell Signaling | 4902S |
| Rabbit anti-PIP5K1A antibody | Cell Signaling | 9693S |
| Rabbit polyclonal anti-phospho-NMDAR (Ser896) antibody | Invitrogen | PA5-37589 |
| Rabbit anti-STIM1 antibody | Cell Signaling | 4916S |
| Rabbit anti-STIM2 antibody | Cell Signaling | 4917S |
| Rabbit polyclonal anti-TMEM24 antibody | Fitzgerald | 70R-7325 |
| Mouse monoclonal anti-TOMM20 antibody | Santa Cruz | sc-11415 |
| Rabbit monoclonal anti-TRKB antibody | Cell Signaling | 4603 |
| Rabbit polyclonal anti-VAP-33 (VAPA) antibody | Bethyl | A304-366A |
| Rabbit polyclonal anti VAPB antibody | Bethyl | A302-894A |
| Cholera toxin subunit B Biotin-conjugated | Invitrogen | C34779 |
| Cholera toxin subunit B Alexa Fluor™ 555-conjugated | Invitrogen | C34776 |
| Cholera toxin subunit B HRP-conjugated | Invitrogen | C34780 |
| Donkey anti-Rabbit Alexa Fluor™ 488-conjugated IgG | Invitrogen | A21206 |
| Bacterial and virus strains |  |  |
| GFP-MAPPER | Addgene | Plasmid #117721 |
| pGP-CMV-GCAMP6s | Addgene | Plasmid #40753 |
| Chemicals, peptides, and recombinant proteins |  |  |
| GM1 isolated from bovine brain | Millipore Sigma | G7641 |
| Protein A Magnetic Beads | Cell Signaling | 73778S |
| NanoLINK Strepavidin Magnetic Beads | Vector Labs | M-1002-010 |
| Critical commercial assays |  |  |
| FD Rapid GolgiStain™ Kit | FD NeuroTechnologies | PK401 |
| Deposited data |  |  |
| ER-PM junction proteomics data | ProteomeXchange | PXD042994 |
| Experimental models: Cell lines |  |  |
| HeLa | ATCC | CRM-CCL-2 |
| HEK293T | ATCC | CRL-3216 |
| Experimental models: Organisms/strains |  |  |
| B6.129P2(FVB)-*Glb1*<sup>tm1Adz/J</sup> (β-*Gal*<sup>-/-</sup>) | Jackson Labs | Strain #:037063 |
| Software and algorithms |  |  |
| Imaris Imaging Software | Imaris |  |
| Other |  |  |
| Pansorbin Cells | Millipore Sigma | 507858-1GM |

## Notes

https://proteomecentral.proteomexchange.org/cgi/GetDataset?ID=PXD042994

### Competing Interest Statement

The authors have declared no competing interest.

