## Supplemental Figures for "Altered GM1 catabolism affects NMDAR-mediated Ca^2+^ signaling at ER-PM junctions and increases synaptic spine formation"

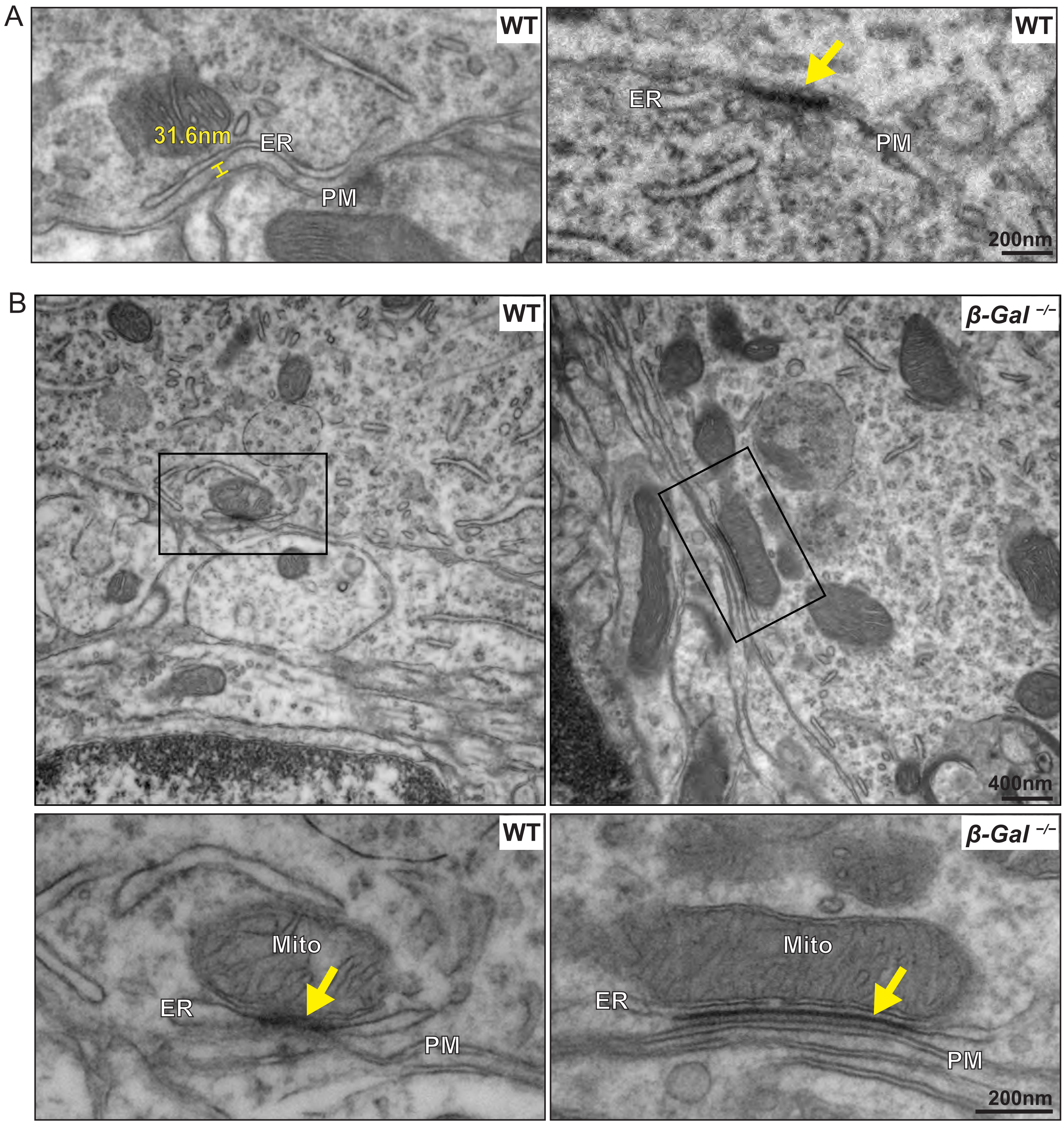


**Figure S1: Purkinje cells form triple-contact sites that connect the ER, PM, and mitochondria. (A)** High-magnification TEM images of Purkinje cells from 6-month-old WT mice show ER-PM junctions defined by the distance between the membranes: the left image shows separation by a distance of ≤35 nm, and the right image shows close apposition resulting in density (yellow arrow). Scale bar: 200 nm. **(B)** TEM images showing triple-contact sites connecting the ER, PM, and mitochondria in WT and β-*Gal*$^{-/-}$ Purkinje cells. Scale bar; 400 nm. Bottom images show 2.5× zoom to magnify the regions in the black boxes in top figure. Scale bar: 200 nm.

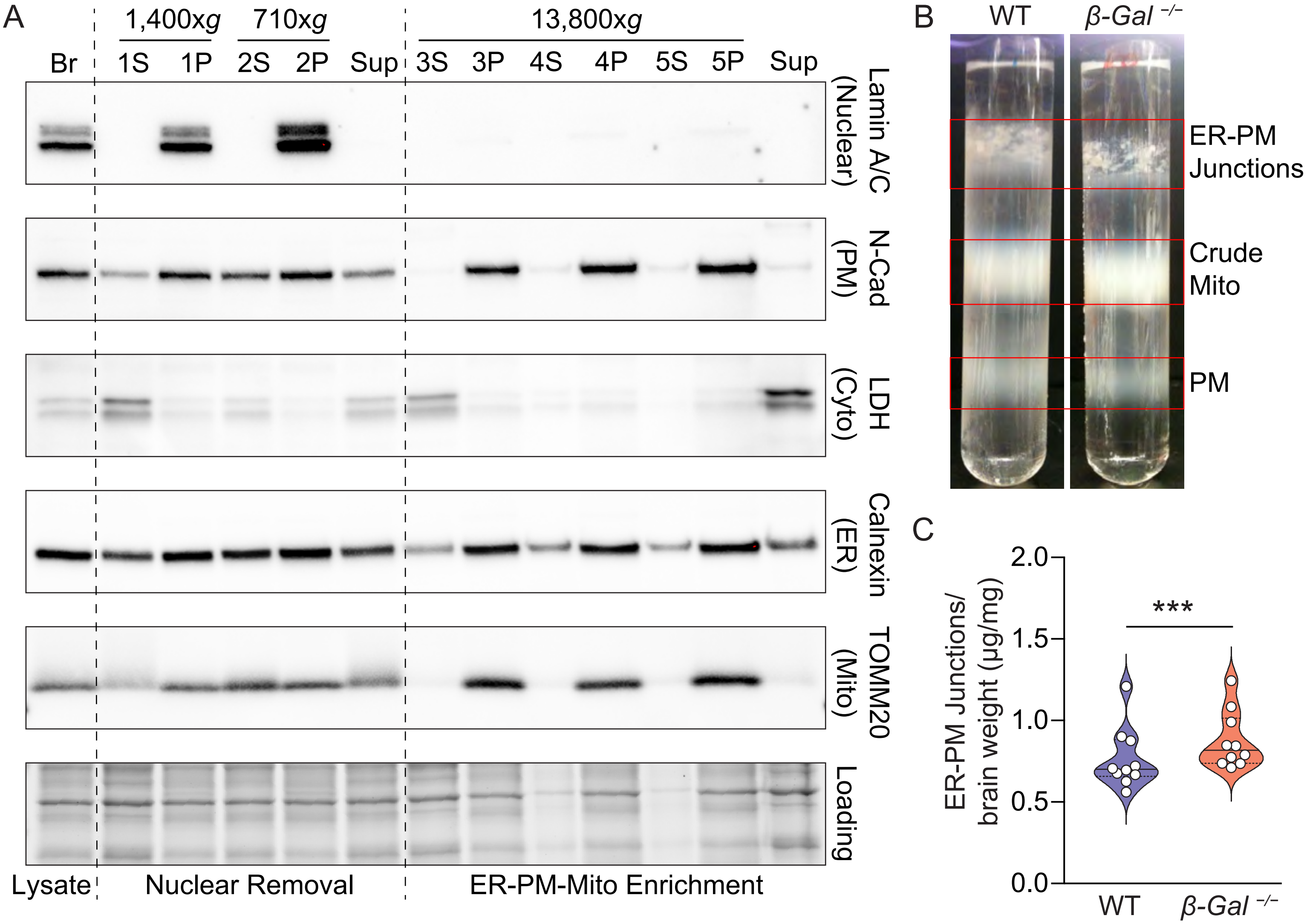


**Figure S2: Purification and validation of ER-PM junctions from WT and β-Gal$^{-/-}$ brains. (A)** Immunoblots showing purity markers of each fraction collected during differential centrifugation of WT and β-*Gal*$^{-/-}$ mouse brains including markers for nucleus (Laminin A/C), ER (Calnexin), PM (N-cadherin), mitochondria (TOMM20), and cytosol (lactose dehydrogenase [LDH]). **(B)** Representative image of the gradient used to separate ER-PM junctions, crude mitochondria, and PM fractions from the 5P pellet in A. **(C)** Quantification of the protein content in ER-PM junctions per brain weight of WT and β-*Gal*$^{-/-}$ mice. Values are expressed as the mean ± SD. Statistical analysis was performed using the Student's t-test; ***$p < 0.001$.

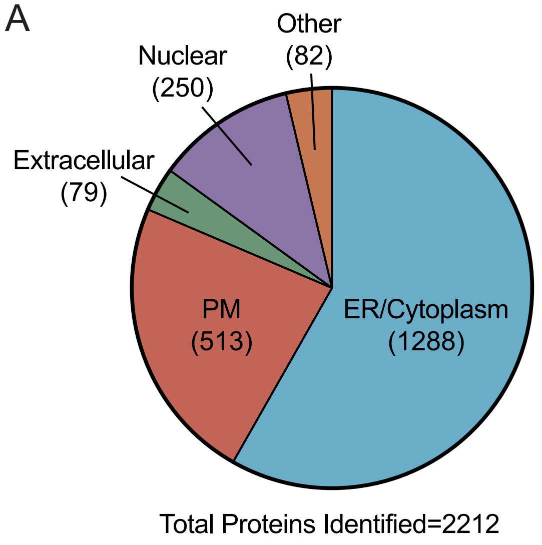


B

| Symbol | Entrez Gene Name | Fold Change |
|---|---|---|
| TMEM24 | Transmembrane protein 192 | 1.30 |
| ORAI2 | ORAI calcium release-activated calcium modulator 2 | 2.00 |
| RYR2 | Ryanodine receptor 2 | 2.22 |
| RYR3 | Ryanodine receptor 3 | 2.00 |
| ITPR3 | Inositol 1,4,5-trisphosphate receptor type 3 | 1.10 |
| GRIN1 | Glutamate ionotropic receptor NMDAR type 1 subunit | 10.00 |
| NTKR2 | Tyrosine Kinase Receptor B | 1.56 |
| MAPK1 | Mitogen-activated protein kinase 1 | 1.33 |
| MAPK3 | Mitogen-activated protein kinase 3 | 1.46 |
| PI4K2A | Phosphatidylinositol 4-kinase type 2 alpha | 20.00 |
| PIP5K1C | Phosphatidylinositol-4-phosphate 5-kinase type 1 gamma | 1.29 |

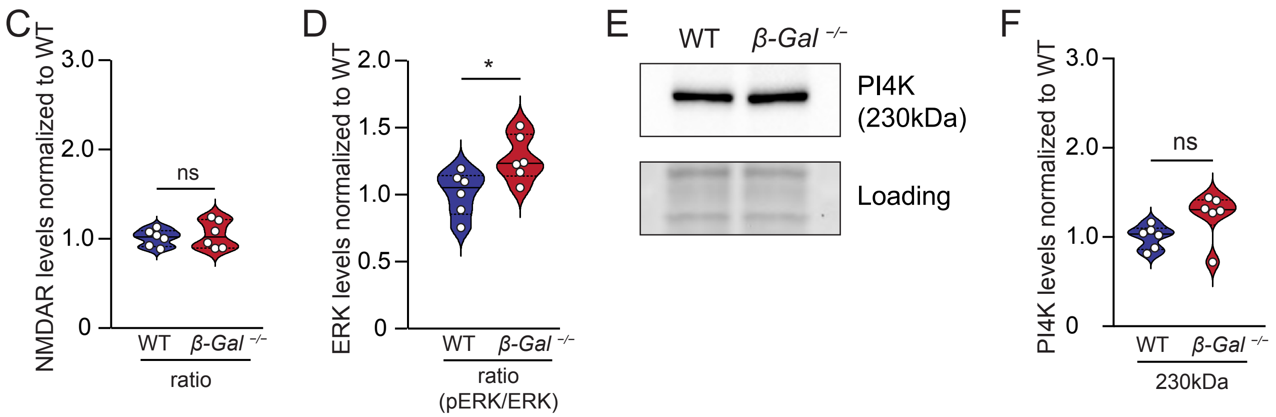


**Figure S3: Ingenuity pathway analysis of proteins identified by proteomics of WT and β-Gal$^{-/-}$ ER-PM junctions. (A)** Pie chart showing the subcellular localization of the proteins identified by the proteomic analysis of WT and β-*Gal*$^{-/-}$ ER-PM junctions. **(B)** Proteomic analysis revealed the upregulation of bone fide ER-PM junction markers (TMEM24 and ORAI2), ER Ca2+-release channels (RYR and ITPR3), synaptic proteins (GRIN1 and NTKR2), ERK-signaling proteins (MAPK1 and MAPK2), and kinases acting on PIs (PI4K2A and PIP5K1C) in β-*Gal*$^{-/-}$ ER-PM junctions. **(C)** Quantification of ratio of NMDAR-type 1 subunit to pNMDAR from Figure 4A. $n$ = 6. Values are expressed as the mean ± SD. Statistical analysis was performed using the Student's t-test. **(D)** Quantification of ratio of ERK to pERK from Figure 4B. $n$ = 6. Values are expressed as the mean ± SD. Statistical analysis was performed using the Student's t-test; *$p < 0.05$. **(E)** Representative immunoblot of P14K band at 230kDa. **(F)** Quantification of immunoblot in E. $n$ = 6. Values are expressed as the mean ± SD. Statistical analysis was performed using the Student's t-test.

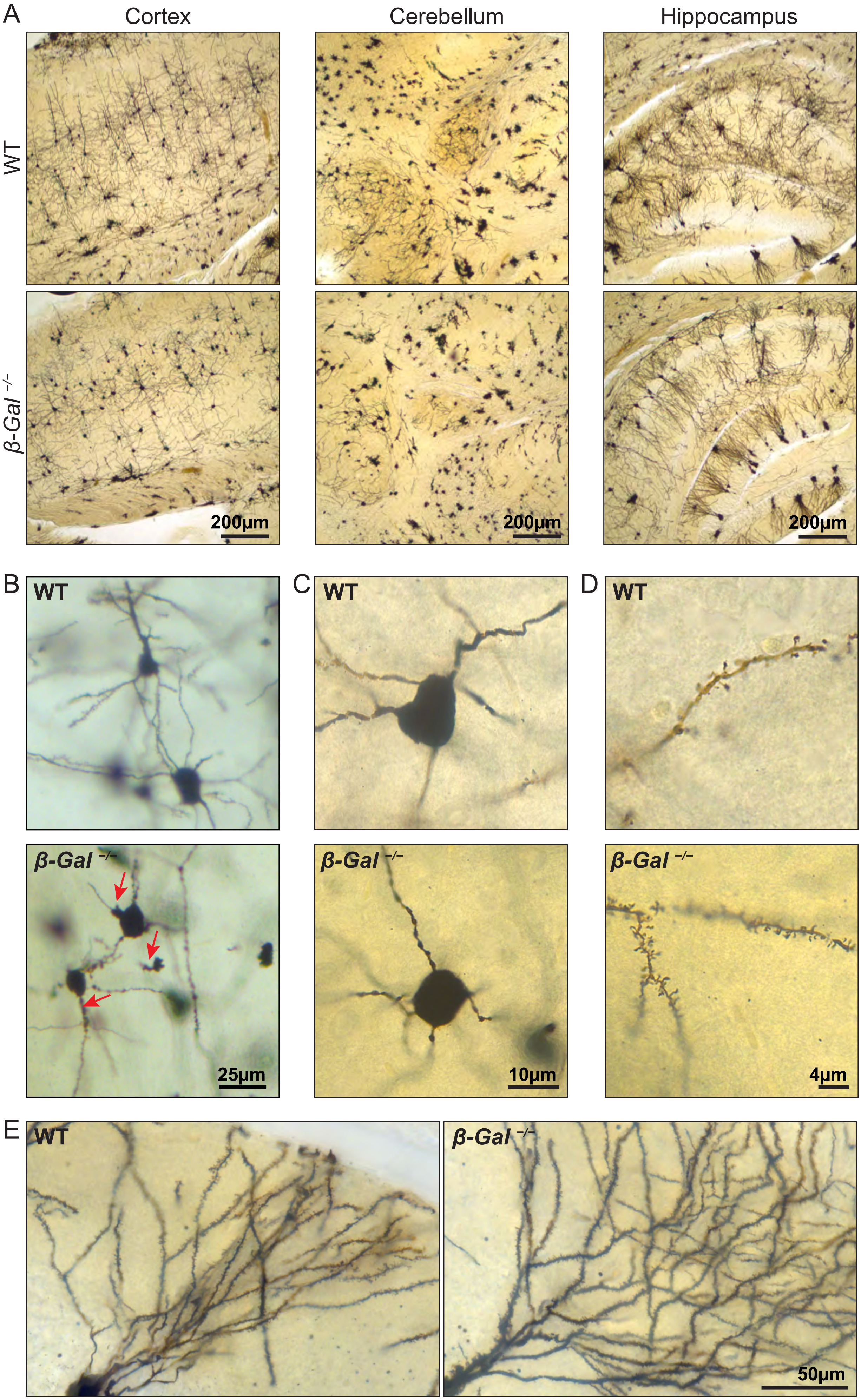

A
Cortex
Cerebellum
Hippocampus
WT
β-Gal −/−
200μm
200μm
200μm
B
C
D
WT
WT
WT
β-Gal −/−
β-Gal −/−
β-Gal −/−
25μm
10μm
4μm
E
WT
β-Gal −/−
50μm

**Figure S4: Golgi-Cox staining of WT and β-*Gal*$^{-/-}$ brains shows dendritic abnormalities in *β-Gal*$^{-/-}$ mice. (A)** Golgi-Cox staining of 6-month-old WT and β-*Gal*$^{-/-}$ showing the cortex, deep cerebellar nuclei region of the cerebellum, and Cornus Ammonis 2 (CA2) region of the hippocampus. Scale bar: 200 µm. **(B)** High-magnification images of Golgi-Cox–stained dendritic arbors in 6-month-old WT and β-*Gal*$^{-/-}$ cortical neurons show the formation of ectopic dendritic growth in β-*Gal*$^{-/-}$ neurons (red arrows). Scale bar: 25 µm. **(C)** High-magnification images of Golgi-Cox–stained dendritic arbors in 6-month-old WT and β-*Gal*$^{-/-}$ cortical neurons show dendritic beading and retracted neural processes in β-*Gal*$^{-/-}$ neurons. Scale bar: 10 µm. **(D)** High-magnification images of Golgi-Cox–stained dendritic arbors in 6-month-old WT and β-*Gal*$^{-/-}$ cortical neurons show increased dendritic spine density in β-*Gal*$^{-/-}$ neurons. Scale bar: 4 µm. **(E)** Golgi-Cox staining of 6-month-old WT and β-*Gal*$^{-/-}$ hippocampal pyramidal cells shows extensive spine formation in β-*Gal*$^{-/-}$ neurons. Scale bar: 50 µm.

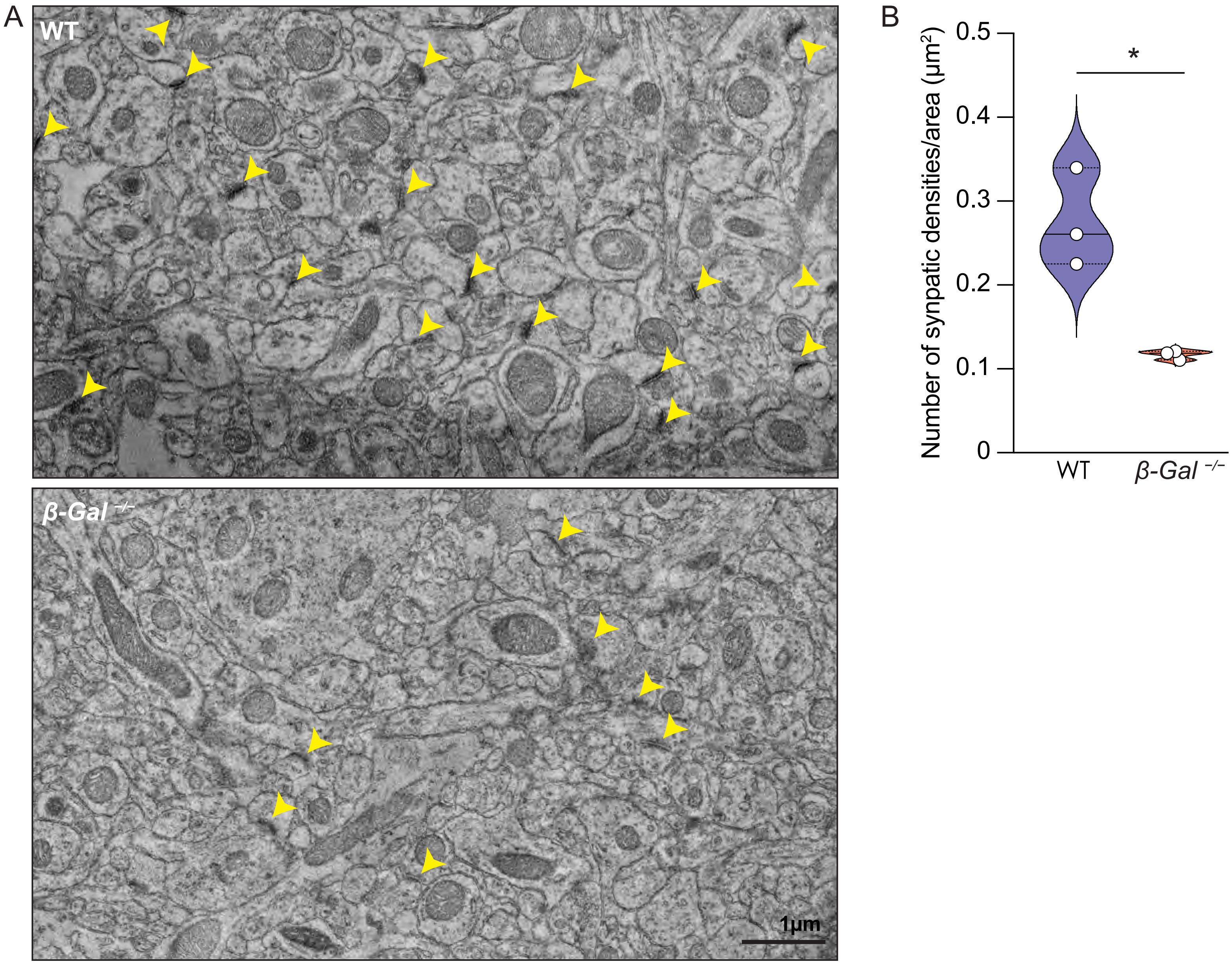


**Figure S5: β-Gal$^{-/-}$ hippocampal pyramidal neurons have fewer synaptic densities than WT neurons. (A)** TEM images of 6-month-old WT and β-*Gal*$^{-/-}$ hippocampal pyramidal cell dendritic arbors. Synaptic densities are labeled by yellow arrows. Scale bar: 1 µm. **(B)** Quantification of the synaptic densities per field of view of the dendritic arbors in A. *n* = 3; 3 mice, 12-15 fields of view along dendritic arbors of CA2 pyramidal cells per mouse. Values are expressed as mean ± SD. Statistical analysis was performed using the Student's t-test with Welch's correction; $^{*}p < 0.05$.

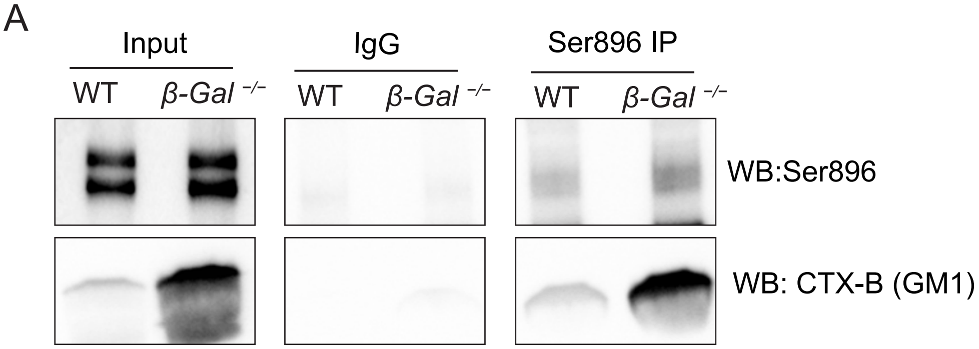


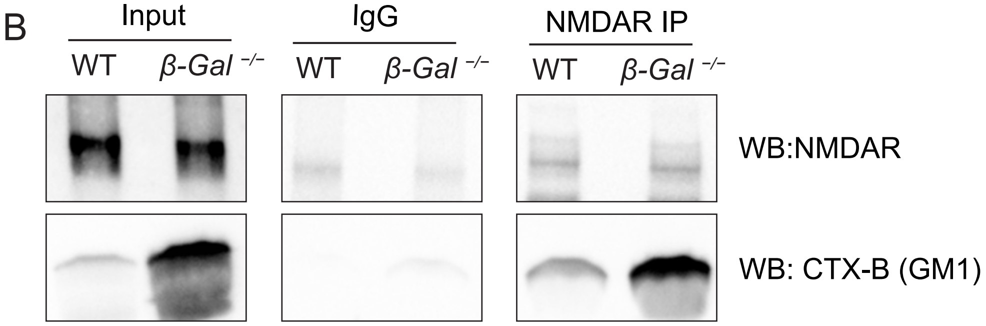


**Figure S6: GM1 binds NMDAR and pNMDAR at the ER-PM junctions in β-Gal$^{-/-}$ mice. (A)** Immunoblot of co-IP using anti-pNMDAR (Ser896) antibody to pull down GM1 in ER-PM junctions isolated from 6-month-old WT and β-*Gal*$^{-/-}$ mice. CTX-B was used to measure GM1 levels. **(B)** Immunoblot of co-IP using anti-NMDAR antibody to pull down bound molecules in ER-PM junctions isolated from 6-month-old WT and β-*Gal*$^{-/-}$ mice. CTX-B was used to measure GM1 levels.

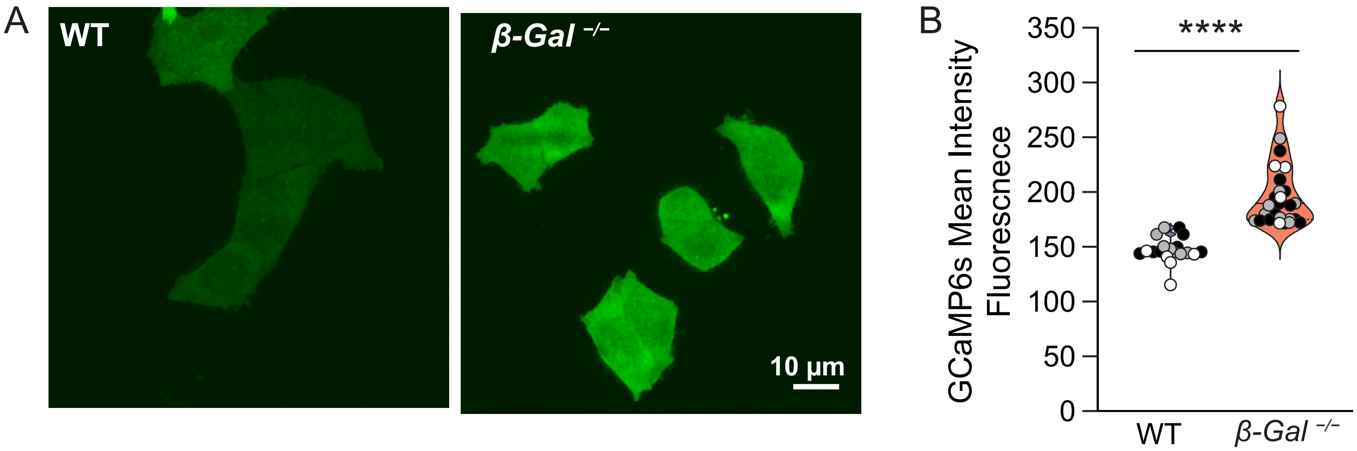


**Figure 7: HeLa cells loaded with GM1 have higher intracellular Ca2+ levels than do control cells. (A)** Fluorescence live-cell Ca2+ imaging using the genetically encoded Ca2+ sensor, GCamp6s, transduced in HeLa cells loaded with GM1. Scale bar: 10 μm. **(B)** Quantification of the intensity of GCamp6s fluorescence in HeLa cells from A. n = 17; 3 passages with 5-6 wells; passage is represented by a different colored filled datapoint (black, grey, white). Values are expressed as mean ± SD. Statistical analysis was performed using the Student's t-test; ****$p < 0.0001$.
